# Tetracistronic Minigenomes Elucidate a Functional Promoter for Ghana Virus and Unveils Cedar Virus Replicase Promiscuity for all Henipaviruses

**DOI:** 10.1101/2024.04.16.589704

**Authors:** Griffin D. Haas, Katharina S. Schmitz, Kristopher D. Azarm, Kendra N. Johnson, William R. Klain, Alexander N. Freiberg, Robert M. Cox, Richard K. Plemper, Benhur Lee

## Abstract

Batborne henipaviruses, such as Nipah virus and Hendra virus, represent a major threat to global health due to their propensity for spillover, severe pathogenicity, and high mortality rate in human hosts. Coupled with the absence of approved vaccines or therapeutics, work with the prototypical species and uncharacterized, emergent species is restricted to high biocontainment facilities. There is a scarcity of such specialized spaces for research, and often the scope and capacity of research which can be conducted at BSL-4 is limited. Therefore, there is a pressing need for innovative life-cycle modeling systems to enable comprehensive research within lower biocontainment settings. This work showcases tetracistronic, transcription and replication competent minigenomes for Nipah virus, Hendra virus, Cedar virus, and Ghana virus, which encode viral proteins facilitating budding, fusion, and receptor binding. We validate the functionality of all encoded viral proteins and demonstrate a variety of applications to interrogate the viral life cycle. Notably, we found that the Cedar virus replicase exhibits remarkable promiscuity, efficiently rescuing minigenomes from all tested henipaviruses. We also apply this technology to GhV, an emergent species which has so far not been isolated in culture. We demonstrate that the reported sequence of GhV is incomplete, but that this missing sequence can be substituted with analogous sequences from other henipaviruses. Use of our GhV system establishes the functionality of the GhV replicase and identifies two antivirals which are highly efficacious against the GhV polymerase.

**Author Summary:** Henipaviruses, such as the prototypical Nipah virus and Hendra virus, are recognized as significant global health threats due to their high mortality rates and lack of effective vaccines or therapeutics. Due to the requirement for high biocontainment facilities, the scope of research which may be conducted on henipaviruses is limited. To address this challenge, we developed innovative tetracistronic, transcription and replication competent minigenomes for Nipah virus, Hendra virus, Cedar virus, as well as for the emergent species, Ghana virus. We demonstrate that these systems replicate key aspects of the viral life cycle, such as budding, fusion, and receptor binding, and are safe for use in lower biocontainment settings. Importantly, application of this system to Ghana virus revealed that its known sequence is incomplete; however, substituting the missing sequences with those from other henipaviruses allowed us to overcome this challenge. We demonstrate that the Ghana virus replicative machinery is functional and identify two orally-efficacious antivirals effective against it. Further, we compare the compatibility of divergent henipavirus replicases with heterotypic viral genetic elements, providing valuable insights for how these species have evolved. Our research offers a versatile system for life-cycle modeling of highly pathogenic henipaviruses at low biocontainment.

## Introduction

Diverse paramyxoviruses circulate in wildlife reservoirs across the globe, with bats hosting a plethora of species(1–4). Bat-borne members of the genus *Henipavirus* pose a major threat to global health, as the prototypical members, Nipah virus (NiV) and Hendra virus (HeV), have demonstrated a propensity for spillover and cause severe encephalitic and respiratory disease in humans(5–7). Furthermore, there have been documented instances of human-to-human transmission of NiV, positioning it as a pathogen of potential pandemic concern(8). As there are no approved vaccines nor therapeutics to prevent or treat henipaviral disease, both NiV and HeV have been designated as select agents and their handling is restricted to high biocontainment facilities (BSL-4). In addition to NiV and HeV, Cedar virus (CedV), a nonpathogenic species, was identified in 2012 and was isolated from *Pteropus* bats in Australia(9–11). There is mounting evidence that henipaviruses (HNVs) circulate within wildlife elsewhere across the globe. Serological studies, for instance, have suggested that henipa-like viruses have spilled over into individuals who were reported to be engaged in bushmeat hunting practices in Cameroon(12). Likewise, metagenomic endeavors have identified an ever-expanding list of new bat-borne paramyxoviruses; these include GhV, sequenced from *Eidolon helvum* bats in Ghana, and Angavokely virus, sequenced from *Eidolon dupreanum* bats in Madagascar(1, 13). Despite these discoveries, successful isolation of HNV species remains limited to NiV, HeV, and CedV; consequently, there are major knowledge gaps in our understanding of emergent, never-before-isolated viruses(11, 14, 15). Until the pathogenicity of newly identified species is assessed, their handling must likewise be restricted to high biocontainment facilities.

The necessary restriction of HNV research with authentic virus to BSL-4 limits the capacity and scope of research which may be conducted, and there is a major need for life cycle modeling systems through which to safely interrogate the viral life cycle of HNVs at lower biocontainment. Monocistronic minigenome systems have been historically employed for NiV and provide a platform through which to safely assess viral RNA dependent RNA polymerase (vRdRp) activity(16, 17); however, these systems are limited in application as they do not provide a means for studying other viral biological processes, such as cytopathic effect (CPE), virus-host interactions beyond the replicase, viral egress, or viral entry. These limitations have been overcome for other viral families with highly pathogenic members, such as *Filoviridae*, through the development of multicistronic minigenomes which encode viral proteins which facilitate diverse aspects of the viral life cycle, but lack the essential viral replicase genes(18).

Henipaviruses, like all paramyxoviruses, follow a strict adherence to the ‘rule of six’, a phenomenon in which genome sizes must be equally divisible by six in length for efficient replication(17, 19). Each monomer of the viral nucleocapsid (N) binds precisely to six nucleotides of the viral RNA, referred to as a ‘hexamer’, and oligomerization of N along the entire length of the vRNA drives a helical, three-dimensional arrangement of the viral ribonucleoprotein (vRNP) complex(20, 21). This helical arrangement facilitates proper phasing of two bipartite promoter elements onto the same surface of the vRNP, facilitating efficient recognition by the viral RNA dependent RNA polymerase (vRdRp). For NiV, promoter element I (PrE-I) is reported to encompass hexamers 1 through 3 (the first 18 nucleotides relative to the 3’ terminal end of the vRNA) and promoter element II (PrE-II) is localized within hexamers 14 through 16 (nucleotides 79 through 96 relative to the 3’ terminal end)(22). Upon recognition of the viral antigenomic promoter, the vRdRp switches exclusively to a replicase mode. In contrast, recognition of the genomic promoter can trigger either replication or can prompt the vRdRp to scan for elements needed to induce transcription of viral mRNAs. In scanning mode, the vRdRp must detect the respective gene start (GS) and gene end (GE) sequences flanking virally-encoded genes, which regulate transcriptional initiation and termination, respectively (23, 24). Violation of the rule of six and/or improper bipartite promoter phasing can significantly hinder the *de novo* rescue of paramyxoviruses by reverse genetics systems. Thus, the biological rules governing paramyxovirus replication are important considerations in the design of reverse genetics systems(25, 26).

Here we report the development of tetracistronic, transcription and replication competent (TC-tr) HNV minigenomes for more versatile life-cycle modeling applications at BSL-2. These systems encode the viral proteins required for budding (M), fusion (F), and receptor binding (RBP) in addition to a reporter gene. We demonstrate that this system is biologically contained, and that all encoded viral proteins are functionally competent. We apply this technology to GhV, an emergent HNV which has never been isolated in culture and use TC-tr minigenomes to explore promoter recognition by the GhV replicase and pinpoint incompatibilities which hinder heterotypic cross-rescue between diverse HNV species. Further, we report for the first time GhV vRdRp susceptibility to two antiviral compounds, contributing to pandemic preparedness efforts in the case GhV, or similar emergent HNVs, spill over into humans.

## Results

### Establishment of a tetracistronic, transcription and replication competent (TC-tr) minigenome for multiple henipavirus species

We constructed tetracistronic, transcription and replication-competent (TC-tr) minigenomes for NiV, HeV, and CedV. This was achieved by cloning the genes responsible for budding (-M), fusion (-F), and receptor binding (-RBP), respectively, within the extreme viral terminal ends (3’ Ldr and 5’ Tr sequences). In addition, we inserted a HiBiT-tagged mCherry reporter gene immediately upstream of HNV-M via a P2A linker, which we have previously demonstrated to have no negative impact on budding in full-length virus (**Fig. 1A and Supplementary Fig. 1A-1C**)(27). To rescue HNV TC-tr minigenomes, BSRT7 cells were co-transfected with the respective TC-tr minigenome plasmids, codon-optimized T7 polymerase, and the cognate accessory plasmids encoding the viral replicase (HNV-N, -P, and -L) (**Fig. 1B and Supplementary Fig. 2**). As a control, each minigenome was rescued in parallel with homotypic HNV-N and -P, but with GFP in lieu of HNV-L. By 48 hours post-transfection, mCherry-positive syncytia were observed for all species, but only when the full viral replicase was provided *in trans*; substitution of the vRdRp (L) with an irrelevant gene (GFP) yielded no rescue (**Fig. 1C**). The rescue efficiency and vRdRp activity of all species was compared by both counting mCherry events per well (**Fig. 1D**) and quantifying relative HiBiT-mCherry expression as a readout for vRdRp activity via nanoluciferase assay (**Fig. 1E**). Rescue of all species yielded thousands of rescue events per well in the presence of HNV-L, with rNiV and rHeV TC-tr minigenomes producing the highest number of mCherry-positive cells (**Fig. 1D**). The HiBiT assay yielded significantly higher RLUs from cells transfected with HNV-L than from cells lacking HNV-L (**Fig. 1E**), further confirming biological containment of the system with a dependency on the vRdRp expressed *in trans*.

**Figure 1.**
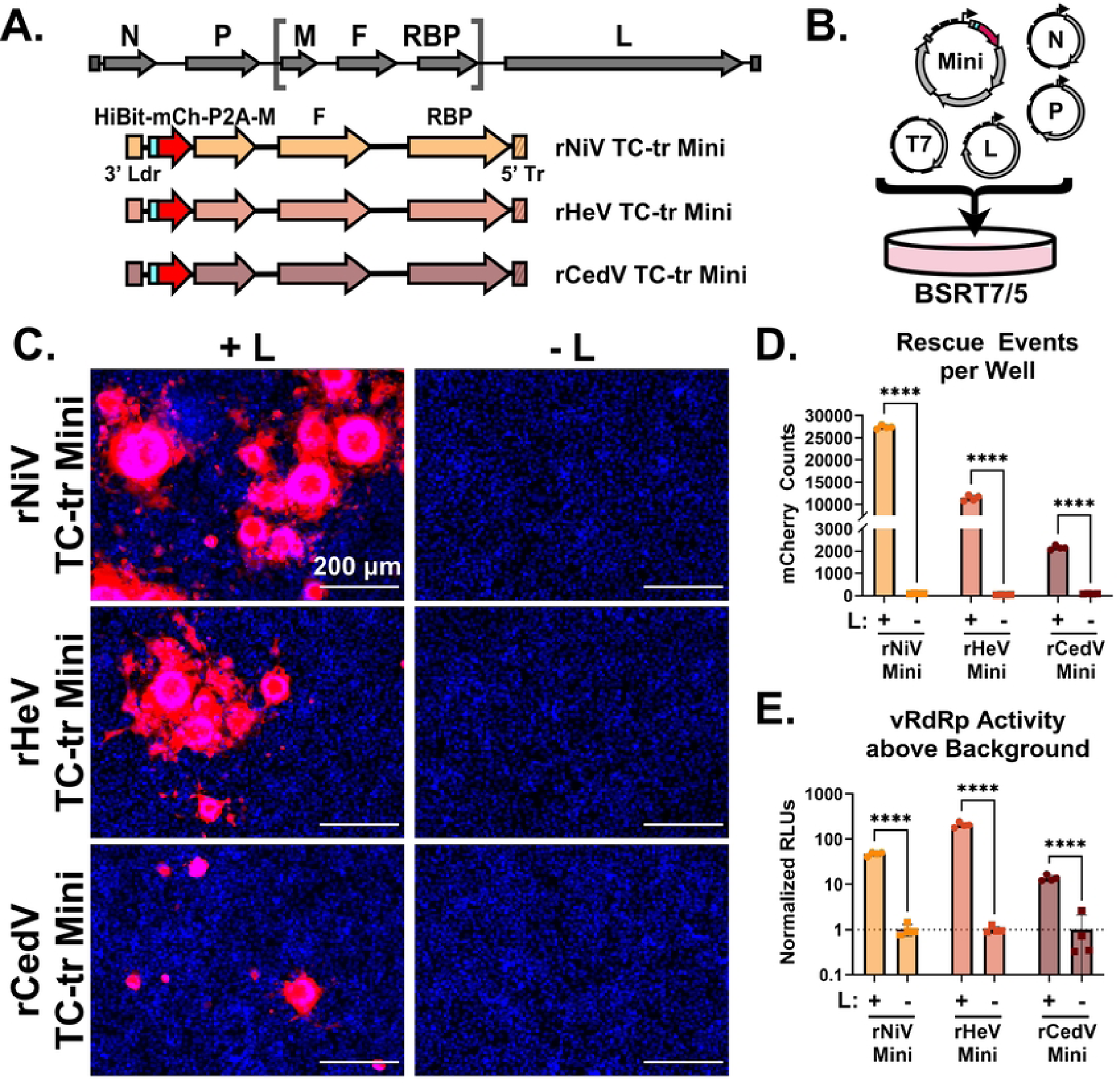
Establishment and rescue of tetracistronic, transcription and replication competent (TC-tr) minigenomes. **(A)** Design of rHNV TC-tr minigenomes which encode a HiBiT-mCherry reporter gene in addition to HNV-M, -F, and -RBP. Full-length virus genome structure is shown above for reference. Detailed descriptions of each minigenome design are available in supplementary figure 1. **(B)** Schematic describing the rescue approach for TC-tr minigenomes in which BSRT7 cells are co-transfected with plasmid encoding the rHNV minigenome, T7-driven HNV-N, -P, -L, and codon-optimized T7 polymerase. A more detailed depiction of the rescue process is available in supplementary figure 2. **(C)** Microscopy demonstrating rescue of rNiV, rHeV, and rCedV TC-tr minigenomes in the presence (+) or absence (−) of their respective HNV-L proteins. For all images, Hoechst is shown in blue while mCherry is shown in red. **(D)** Quantification of mCherry positive events produced by rHNV TC-tr minigenome rescue and **(E)** matched quantification of normalized nanoluciferase signal. To measure vRdRp activity, RLUs from rescues in the presence of HNV-L were normalized to RLUs from rescues in the absence of HNV-L. Statistical significance was assessed by multiple unpaired t tests in GraphPad Prism to compare counts or normalized RLUs in the presence of HNV-L with counts or normalized RLUs in the absence of HNV-L for each respective minigenome. Rescues were conducted in quadruplicate. Error bars depict standard deviation. For all graphs (ns) = P > 0.05; (*) = P ≤ 0.05; (**) = P ≤ 0.01; (***) = P ≤ 0.001; (****) = P ≤ 0.0001.

### Nipah virus matrix protein interacts with nucleolar host factors during TC-tr minigenome infection and is competent for budding

We have previously demonstrated that NiV-M protein is transiently trafficked into nuclei during infection(28). During this sojourn, NiV-M interacts with a broad array of nuclear and nucleolar host factors, including components of the host box C/D snoRNP complex (NOP56, NHP2L1, NOP58, and FBL)(29, 30). Transient nuclear trafficking of HNV-M leads to ubiquitination of HNV-M, which is required for matrix to be competent in its facilitating budding of VLPs(28). To determine if NiV-M expressed under viral promoter by our TC-tr minigenome undergoes comparable trafficking to nucleoli, we prepared constructs for bimolecular fluorescence complementation (BiFC) utilizing a split mVenus fluorescent reporter. Co-expression of NOP56 fused to VC155 with either NHP2L1 or FBL fused to VN173 yielded appropriate localization of BiFC signal to nucleolar punctae in co-transfection studies (**Supplementary Fig. 3A and 3B**). Further, co-transfection of VN173-fused NOP56 with VC155-fused NiV-M likewise reconstituted fluorescent signal in nucleolar punctae (**Supplementary Fig. 3C**). Confirming that BiFC signal is specific to the respective pair of interaction partners, co-transfection of VN173-fused NiV-M with VC155-fused NiV-M resulted in diffuse cytoplasmic expression (**Supplementary Fig. 3D**).

Having validated our BiFC constructs, we encoded VC155-fused NiV-M into our rNiV TC-tr minigenome (**Supplementary Fig. 3E**). rNiV TC-tr minigenomes encoding either untagged (wildtype, WT) NiV-M or NiV-M fused on its N-terminus with VC155 were rescued in BSRT7 cells. In parallel, HeLa cells were transfected with either NHP2L1 or NOP56 fused with VN173 on their C-terminus. At 12 hours post-transfection (HPT), HeLa cells and BSRT7 cells were trypsinized, mixed, and co-cultured for 48 hours. Fusion mediated by NiV-F and -RBP of BSRT7 cells with HeLa cells should result in cytoplasmic mixing and BiFC (**Fig. 2A**). While TC-tr minigenomes encoding untagged NiV-M failed to yield detectable BiFC, minigenomes encoding VC155 fused with NiV-M produced syncytium containing BiFC signal. Importantly, BiFC was localized to punctae within nuclei, recapitulating our transfection results and demonstrating that NiV-M driven by viral promoter is trafficked to nucleoli during infection (**Fig. 2B**). Having observed that NiV-M undergoes appropriate trafficking, we next assessed its competency for budding. Because TC-tr minigenomes encode the necessary and sufficient proteins for particle formation, fusion, and receptor binding (HNV-M, -F, and -RBP), we anticipated that rescue events would produce viral like particles (TC-trVLPs) which may be used to deliver the minigenomic vRNPs to secondary target cells expressing replicase *in trans*. Indeed, RT-qPCR of supernatant from rNiV and rHeV TC-tr rescue cells yielded upwards of 10^6^ genome copies per mL (**Fig. 2C**). BSRT7 cells were pre-transfected with respective HNV -N, -P, and -L and infected with TC-tr VLP rescue supernatant, yielding mCherry-positive events by 48 hours post infection (HPI). These events were quantified at each passage. These TC-tr VLPs could be iteratively passaged up to two times, albeit with incremental loss in titer (**Fig. 2D**).

**Figure 2.**
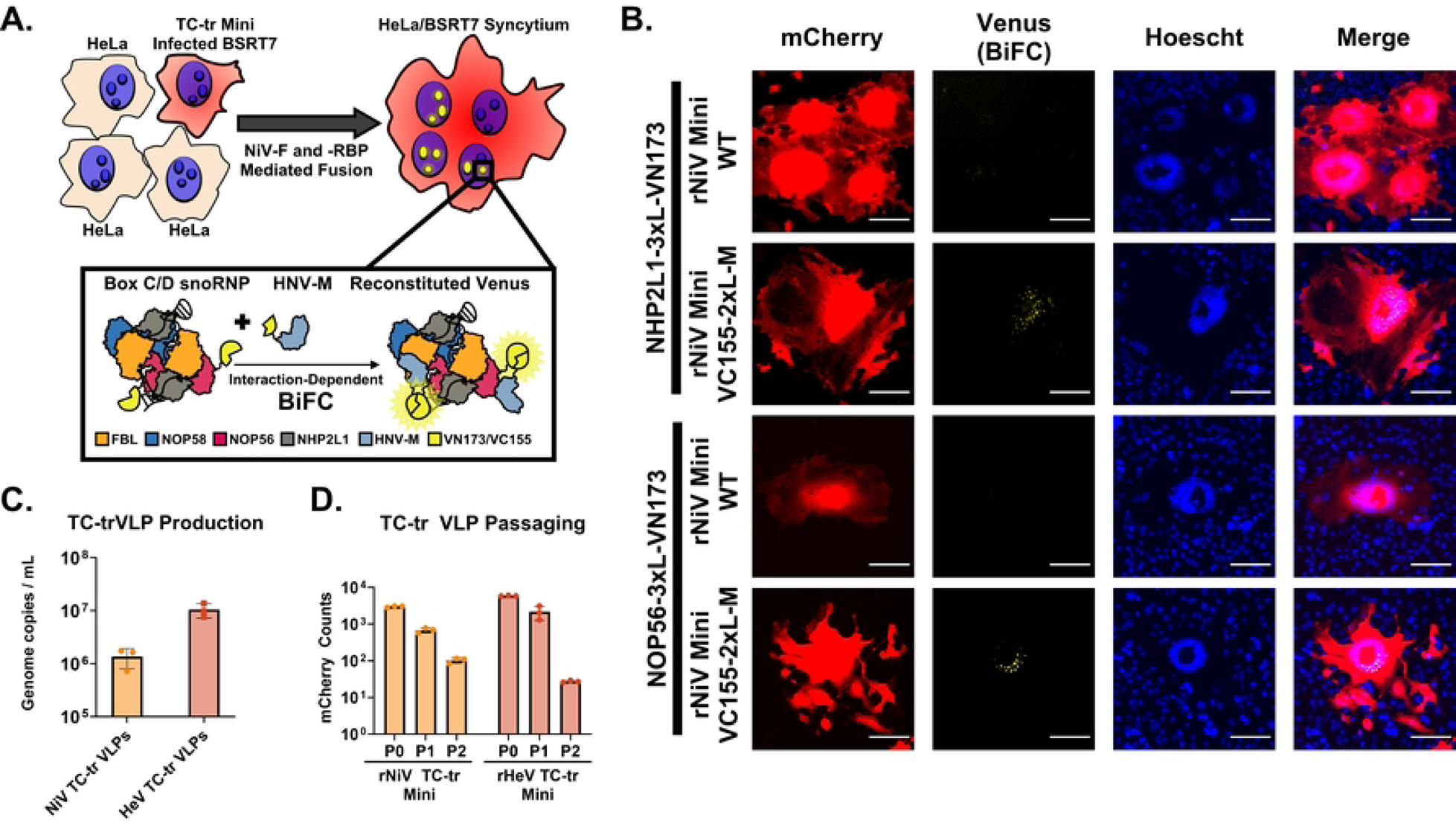
Matrix protein encoded by rHNV TC-tr VLPs is functionally competent. **(A)** Diagram depicting the experimental approach used for the BiFC assay. HeLa cells transfected with VN173-fused NHP2L1 or VN173-fused NOP56 were co-cultured with BSRT7 cells infected with rNiV TC-tr minigenomes encoding WT NiV-M or NiV-M fused with VC155. Fusion mediated by NiV-F and NiV-RBP facilitates cytoplasmic mixing and results in BiFC within mCherry-positive syncytia. Inset cartoon demonstrates reconstitution of Venus fluorescent protein if both VC155-fused NiV-M and VN173-fused host box C/D snoRNP proteins interact. **(B)** Microscopic images capturing instances of BiFC within mCherry-positive syncytia. For all images, mCherry is colored in red, BiFC (venus reconstitution) is colored in yellow, and Hoechst stain is colored in blue. Scale bars represent 100 micrometers. **(C)** Detection of TC-tr minigenome vRNA copies in supernatant from rescue cells. **(D)** Iterative passaging of rNiV and rHeV TC-tr VLPs captured by quantification of mCherry positive events at each passage (P0 = rescue; P1 = passage 1; P2 = passage 2). Rescues and passaging experiments were conducted in triplicate. Error bars depict standard deviation.

### The reported genome sequence of Ghana virus (strain m74a) is missing 28nt from its 3’ Leader sequence

We next sought to develop a TC-tr minigenome for GhV, a HNV species which has never been isolated in culture. Unfortunately, a frequent technical challenge in the sequencing of negative-sense, single-stranded RNA viruses is that the terminal ends are often incompletely mapped(31). To determine if the terminal ends of GhV are complete, we aligned the 3’ Ldr and 5’ Tr sequences, respectively, of NiV, HeV, CedV, and GhV by constraining to proper phasing of the genomic and antigenomic bipartite promoter elements. Proper phasing of the bipartite promoters revealed a conservation of the core PrE-II sequence, consisting of the motif ‘ACC’ between hexamers 13 and 14, 14 and 15, and 15 and 16, respectively (**Fig. 3A and 3B**); this consensus sequence is in agreement with previous characterization of PrE-II for NiV(22). This constrained alignment allowed us to elucidate with confidence that the reported sequence of GhV is missing exactly 28 nucleotides from its 3’ Ldr (**Fig. 3A**) while its 5’ Tr is complete (**Fig. 3B**). Furthermore, this constrained alignment properly positions the N gene start (GS) of GhV to begin at position 56, a phasing which is absolutely conserved among all known paramyxoviruses (**Supplementary Fig. 4**)(20). The incomplete terminal end of GhV is of major consequence because the genomic PrE-I, which is essential for vRdRp recognition, exists in the terminal end of the 3’Ldr sequence. Both the PrE-I and PrE-II sequences must be present and properly phased when the vRNA is fully encapsidated in N to form a functional vRNP (**Fig. 3C and Supplementary Fig. 2**). Attempts to construct a GhV minigenome based solely on the available sequence would yield a minigenome lacking a functional genomic PrE-I sequence, which would not be recognizable by the GhV vRdRp (**Fig. 3D**).

**Figure 3.**
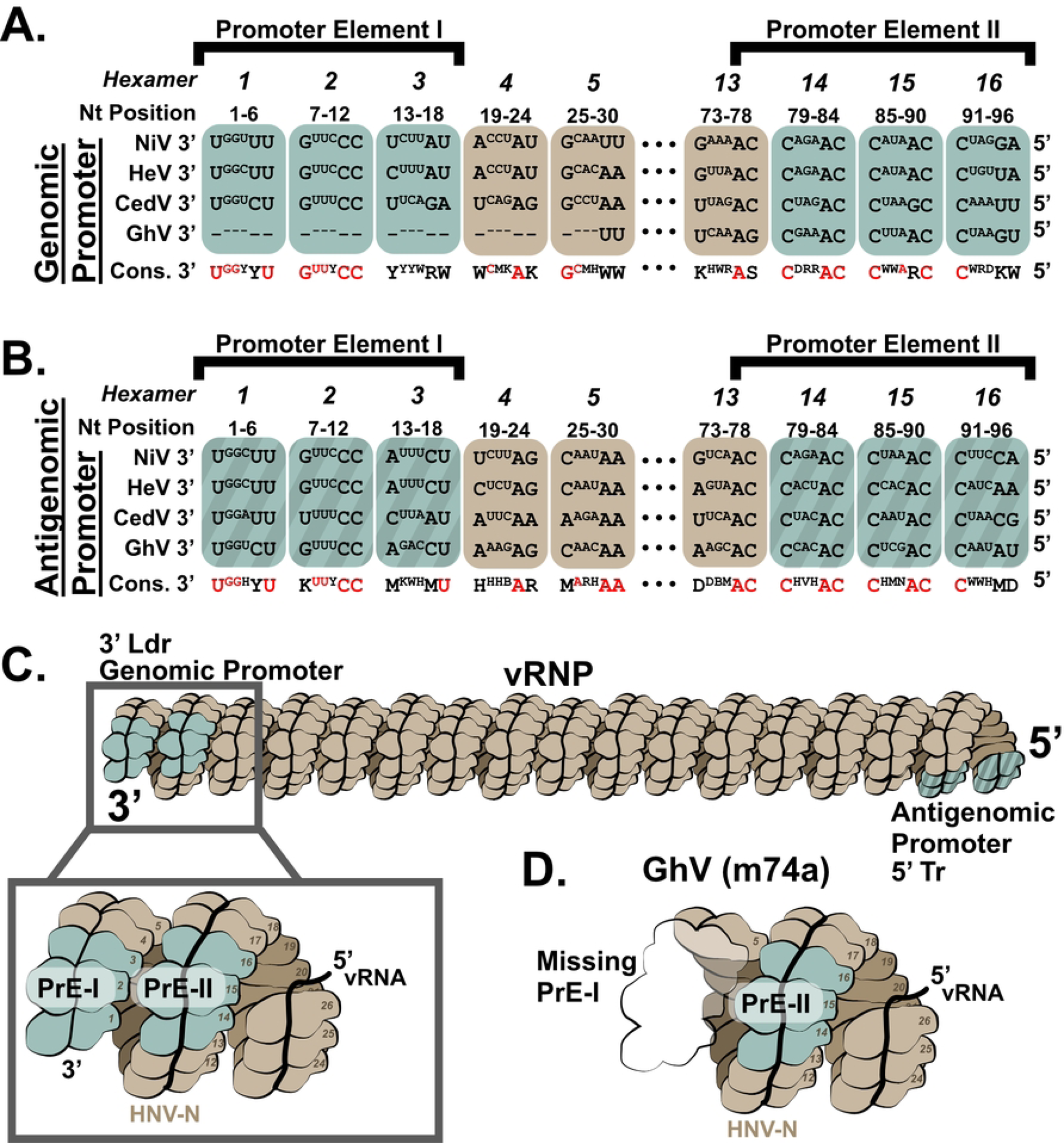
Constrained sequence alignment of the GhV genome reveals that GhV is missing 28 nucleotides from its genomic promoter. **(A)** Sequence alignment of the reported genomic vRNA sequences of NiV, HeV, CedV, and GhV with constraint to proper phasing of the genomic promoter element II (PrE-II) sequence. **(B)** Sequence alignment of the reported antigenomic vRNA sequences of NiV, HeV, CedV, and GhV with constraint to proper phasing of the antigenomic PrE-II sequence. All alignments are shown in a 3’ to 5’ orientation, reflective of the biologically relevant sequences. Bases that are outwards facing towards solvent are shown in full case, while bases buried within the nucleocapsid protein are shown in superscript text. Dashes (−) are used to denote unmapped nucleotides missing in the GhV sequence. The consensus sequence is shown below each hexamer, with absolutely conserved residues colored in red. All nucleotides follow IUPAC nomenclature. **(C)** Cartoon depicting a vRNP with properly phased bipartite promoters on each end. The inset panel shows the relative localization of PrE-I and PrE-II within the genomic promoter, with numbers detailing each hexamer position relative to the 3’ end of the vRNA. This model reflects phasing observed in the cryoEM structure of the NiV helical nucleocapsid assembly (PDB 7NT5). **(D)** Cartoon depicting hexamers and their relative positions in the genomic promoter of the GhV vRNP which correspond to incompletely mapped GhV sequence. Missing hexamers/sequence are shown as a transparent outline. For all figures, monomers of nucleocapsid containing a hexamer of vRNA are denoted in brown, except for hexamers 1-3 and 14-16; hexamers encoding elements of the bipartite promoter elements are colored in teal. The antigenomic bipartite promoter elements are depicted with additional shaded pattern.

### Restoration of a functional genomic promoter is sufficient for the rescue of a rGhV TC-tr minigenome

In an effort to restore a functional genomic PrE-I within the sequence of GhV, we designed rGhV TC-tr minigenomes in which we replaced the missing terminal 28 nucleotides of the GhV 3’ Ldr sequence with cognate sequences derived from the genomic promoters of either NiV (NiV Ldr28), HeV (HeV Ldr28), or CedV (CedV Ldr28), respectively. In addition, a minigenome was constructed in which the terminal 28 nucleotides of the fully-sequenced GhV 5’ Tr (GhV Tr28) was mirrored and encoded in lieu of the missing nucleotides in the GhV 3’ Ldr (**Fig. 4A and Supplementary Fig. 1D**). Co-transfection of BSRT7 cells with the GhV replicase and respective minigenomes resulted in significant rescue only when the NiV Ldr28 or HeV Ldr28 sequences were encoded, while the CedV Ldr28 sequence did not promote rescue above background. (**Fig. 4B and 4C**). Successful rescue of the rGhV minigenome (with HeV Ldr28) yielded modest mCherry-positive events exclusively in the presence of GhV-L (**Fig. 4D**). The GhV Tr28 construct only yielded low levels of rescue above background, suggesting that sequences encoded in the 5’ Tr (antigenomic PrE-I) modulate vRdRp activity differently than do sequences encoded in the 3’ Ldr (genomic PrE-I). Further experimentation confirmed that replacement of the first 28 nucleotides of the 3’ Ldr sequence with the mirrored terminal 28 nucleotides of the viral 5’ Tr (Tr28) resulted in a significant decrease in respective vRdRp activity for both the GhV minigenome (**Fig. 4E**) and the HeV minigenome (**Fig. 4F**) when rescue was performed with their respective replicases.

**Figure 4.**
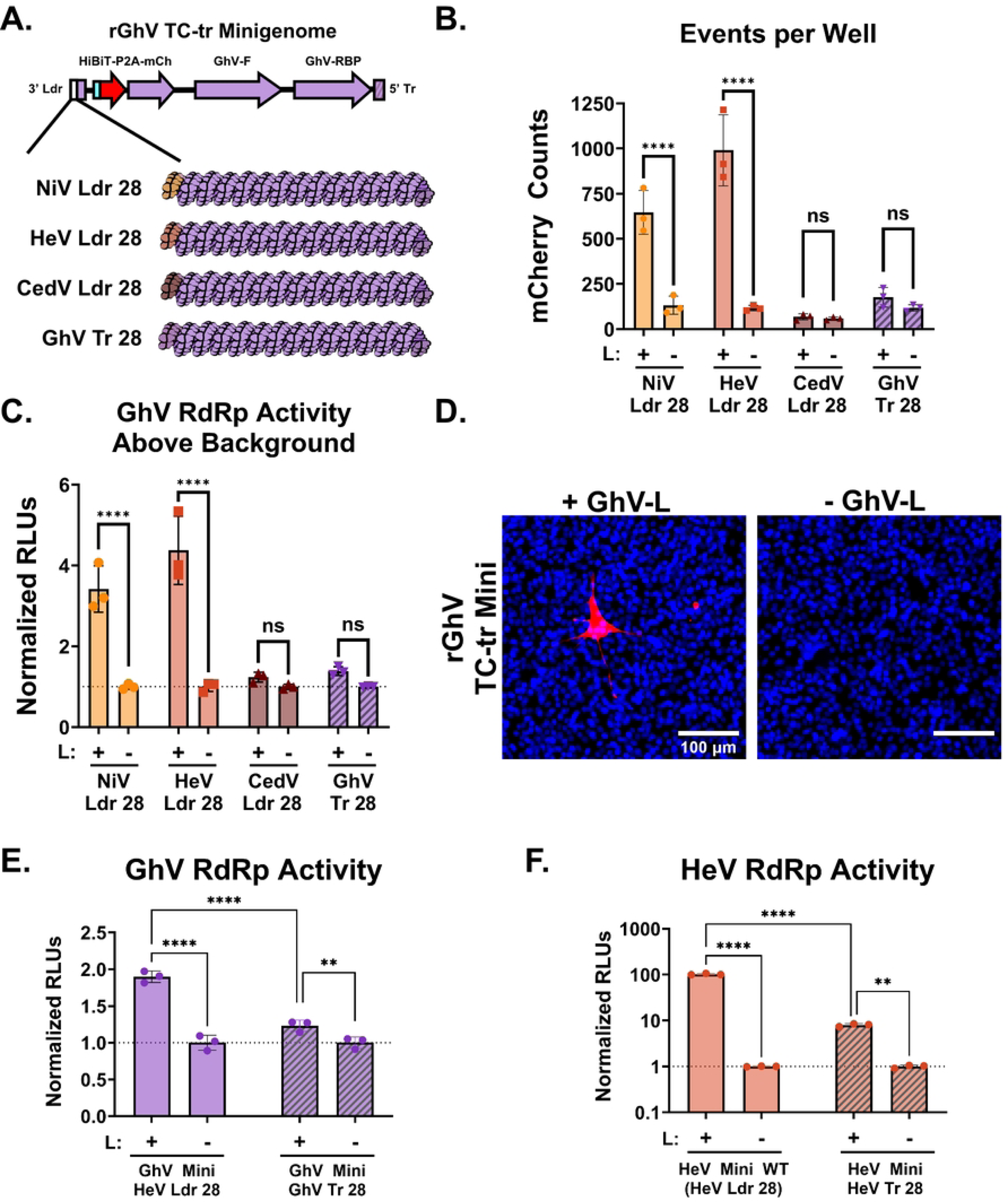
Restoration of a functional PrE-I facilitates rescue of a rGhV TC-tr minigenome. **(A)** Cartoon depicting the design of chimeric rGhV TC-tr minigenomes in which the unmapped terminal 28 nucleotides of the GhV 3’ Ldr sequence are replaced with equivalent sequences derived from NiV (NiV Ldr28), HeV (HeV Ldr28), CedV (CedV Ldr28), or the GhV antigenomic promoter (GhV Tr28). The minigenome design of rGhV is further detailed in supplementary figure 1D. **(B)** Quantification of mCherry positive events and **(C)** matched vRdRp activity resulting from the rescue of each rGhV TC-tr minigenome in the presence (+) or absence (−) of GhV-L. **(D)** Microscopy depicting rGhV (HeV Ldr28) TC-tr minigenome rescue in the presence or absence of GhV-L. Red depicts mCherry signal while blue depicts Hoechst stain. **(E)** Comparison of GhV vRdRp activity resulting from the rescue of rGhV TC-tr minigenomes encoding either the HeV Ldr28 sequence or the GhV Tr28 sequence in the presence or absence of GhV-L. **(F)** Quantification of HeV vRdRp activity resulting from the rescue of rHeV TC-tr minigenomes encoding either the WT (HeV Ldr28) sequence or the HeV Tr28 sequence in the presence or absence of HeV-L. To calculate vRdRp activity, RLUs from rescues in the presence of HNV-L were normalized to respective RLUs generated by each minigenome in the absence of HNV-L. Statistical significance was assessed using a 2way ANOVA analysis in GraphPad Prism to compare counts or normalized RLUs in the presence of HNV-L with respective counts or normalized RLUs in the absence of HNV-L. For **(E)** and **(F)**, additional comparisons were conducted to determine significance between respective minigenome mutants rescued in the presence of HNV-L. All rescues were conducted in triplicate. Error bars depict standard deviation. For all graphs (ns) = P > 0.05; (*) = P ≤ 0.05; (**) = P ≤ 0.01; (***) = P ≤ 0.001; (****) = P ≤ 0.0001.

### The Cedar virus RNA Dependent RNA Polymerase possesses a remarkable plasticity in its template recognition

Having observed that the GhV replicase can recognize the genomic PrE-I sequence derived from both NiV and HeV, but not from CedV (**Fig. 4B and 4C**), we next sought to determine if the replicase of a given HNV species is capable of rescuing the minigenome of heterotypic species. Each TC-tr minigenome was rescued using replicase (N, P, and L) from either NiV, HeV, CedV, or GhV. As a control, each minigenome was rescued in parallel with homotypic HNV-N and -P, but with GFP in lieu of HNV-L. At 72 HPT, nanoluciferase assay was performed to assess vRdRp activity. Curiously, while the NiV replicase and HeV replicase could rescue each other’s minigenomes (**Fig. 5A and 5B**), neither the HeV replicase nor the NiV replicase could rescue a GhV minigenome encoding the HeV Ldr28 sequence (**Fig. 5C**). Likewise, the GhV replicase was only capable of rescuing its homotypic minigenome encoding HeV Ldr28, but not the heterotypic NiV, HeV, nor CedV minigenomes (**Fig. 5A - 5D**). Neither the NiV replicase, HeV replicase, nor GhV replicase was capable of rescuing the rCedV minigenome (**Fig. 5D**). Remarkably, the CedV replicase was capable of rescuing all of the heterotypic HNV minigenomes, demonstrating a marked plasticity in its template recognition relative to the other species (**Fig. 5A-5D**). Heterotypic cross-rescue results are summarized in **Fig. 5E**.

**Figure 5.**
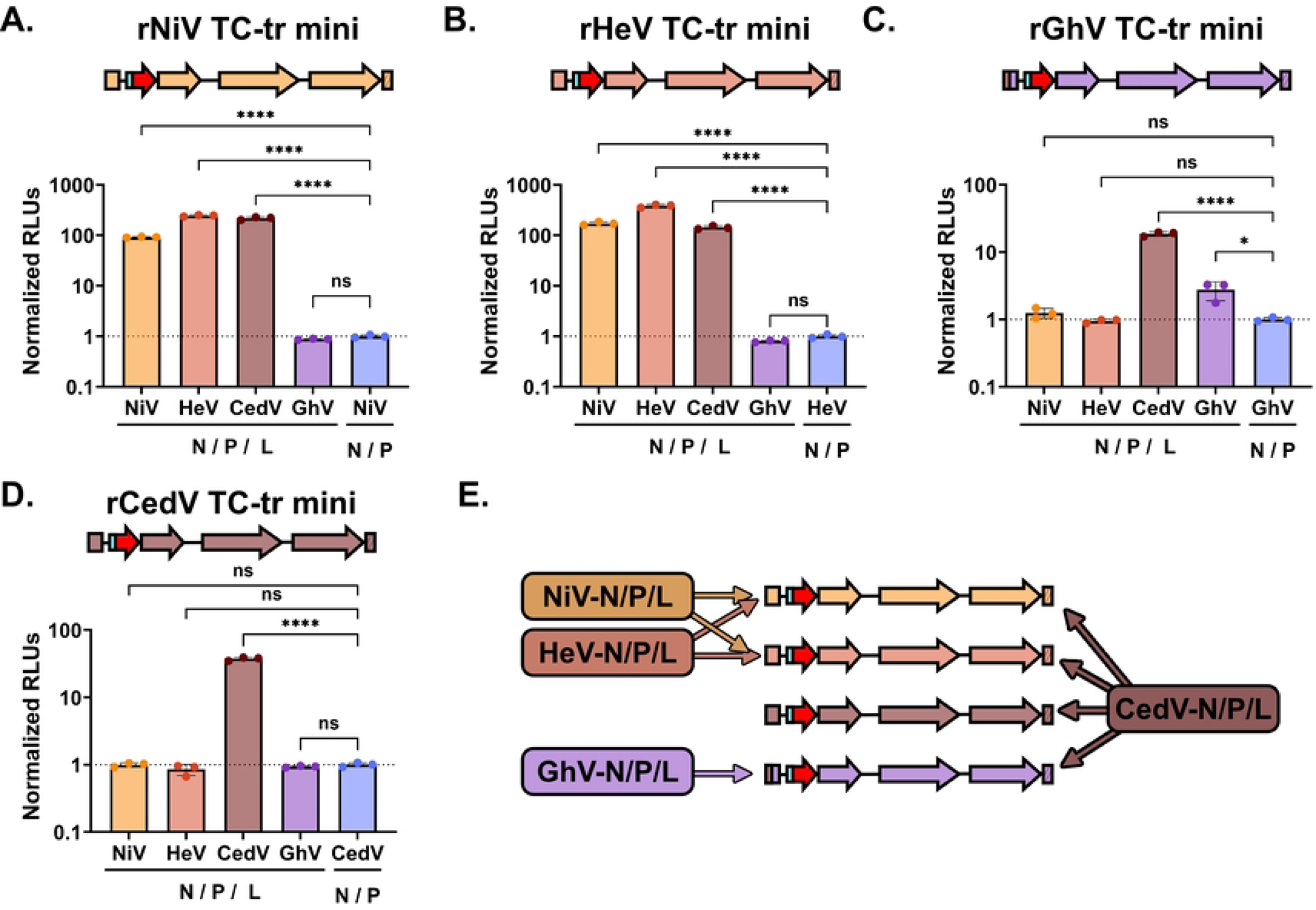
Heterotypic cross-rescue of diverse henipaviruses species uncovers a remarkable plasticity in template recognition by the CedV vRdRp. vRdRp activity above background from the rescue of **(A**) rNiV, **(B**) rHeV, **(C)** rGhV, or **(D)** rCedV TC-tr minigenomes by NiV-N/-P/-L, HeV-N/-P/-L, CedV-N/-P/-L, or GhV-N/-P/-L, respectively. RLUs from each rescue condition were normalized to RLUs from respective minigenome rescues conducted in parallel but in the absence of HNV-L. **(E)** Schematic summarizing the capability of each HNV replicase to successfully rescue each rHNV TC-tr minigenome. Statistical significance was determined by ordinary, one-way ANOVA analysis in GraphPad Prism comparing the normalized RLUs yielded by each HNV replicase species to normalized RLUs from the no-L control. All rescues were conducted in triplicate. Error bars demonstrate standard deviation. For all graphs (ns) = P > 0.05; (*) = P ≤ 0.05; (**) = P ≤ 0.01; (***) = P ≤ 0.001; (****) = P ≤ 0.0001.

### Incompatibilities exist between the respective gene starts, 3’ trailer sequences, and replicases of Hendra virus and Ghana virus

While both the GhV replicase and HeV replicase could successfully recognize the HeV genomic PrE-I (HeV Ldr28 sequence) in the context of their own, homotypic minigenome, neither replicase could support rescue of heterotypic minigenomes encoding the HeV Ldr28 sequence. We hypothesized that there are one or more restrictions in place preventing successful heterotypic cross-rescue. For minigenome rescue to occur there must be (1) sufficient recognition of the antigenomic 5’ Tr sequence by the vRdRp to produce genomic vRNA, (2) recognition of the 3’ Ldr sequence in the genomic vRNA by the vRdRp, and (3) recognition of the first GS sequence by the vRdRp, which is sufficient to engage the polymerase into a transcriptase mode, yielding viral mRNAs and concomitant expression of the HiBiT-mCherry reporter gene (**Fig. 6A and Supplementary Fig. 2**). Because there are differences in the GS sequences controlling reporter gene expression **(Fig. 6B**) between the rGhV and rHeV minigenomes, we constructed a rGhV minigenome in which we replaced the GhV-N GS, which controls reporter gene expression, with the analogous GS sequence from HeV (**Supplementary Fig. 5B**). Likewise, because GhV and HeV have distinct 5’ Tr (antigenomic PrE-I) sequences (**Fig. 3B**), we constructed a rGhV minigenome in which the terminal 28 nucleotides of the 5’ Tr was replaced by the Tr28 sequence from HeV (**Supplementary Fig. 5C**). Finally, we constructed a rGhV minigenome in which both the GS and the Tr28 sequences of GhV were replaced with the analogous sequences from HeV (**Supplementary Fig. 5D**). Rescue of these minigenomes by the GhV replicase demonstrated a clear loss in vRdRp activity when the GS was derived from HeV. Further, the GhV replicase was unable to support the rescue of any minigenome encoding the HeV Tr28 sequence, demonstrating a clear incompatibility between the GhV replicase and HeV antigenomic PrE-I (**Fig. 6C**). Conversely, while the HeV replicase was unable to rescue a rGhV minigenome encoding only the HeV Ldr28, mutating either the GhV-N GS to the HeV-N GS or mutating the GhV Tr28 sequence to the HeV Tr28 sequence was sufficient for some restoration of HeV vRdRp activity. Further, there was a clear, positive combinatorial increase in HeV vRdRp activity when both the GhV-N GS and GhV Tr28 sequences were simultaneously mutated to their respective HeV equivalents (**Fig. 6D**).

**Figure 6.**
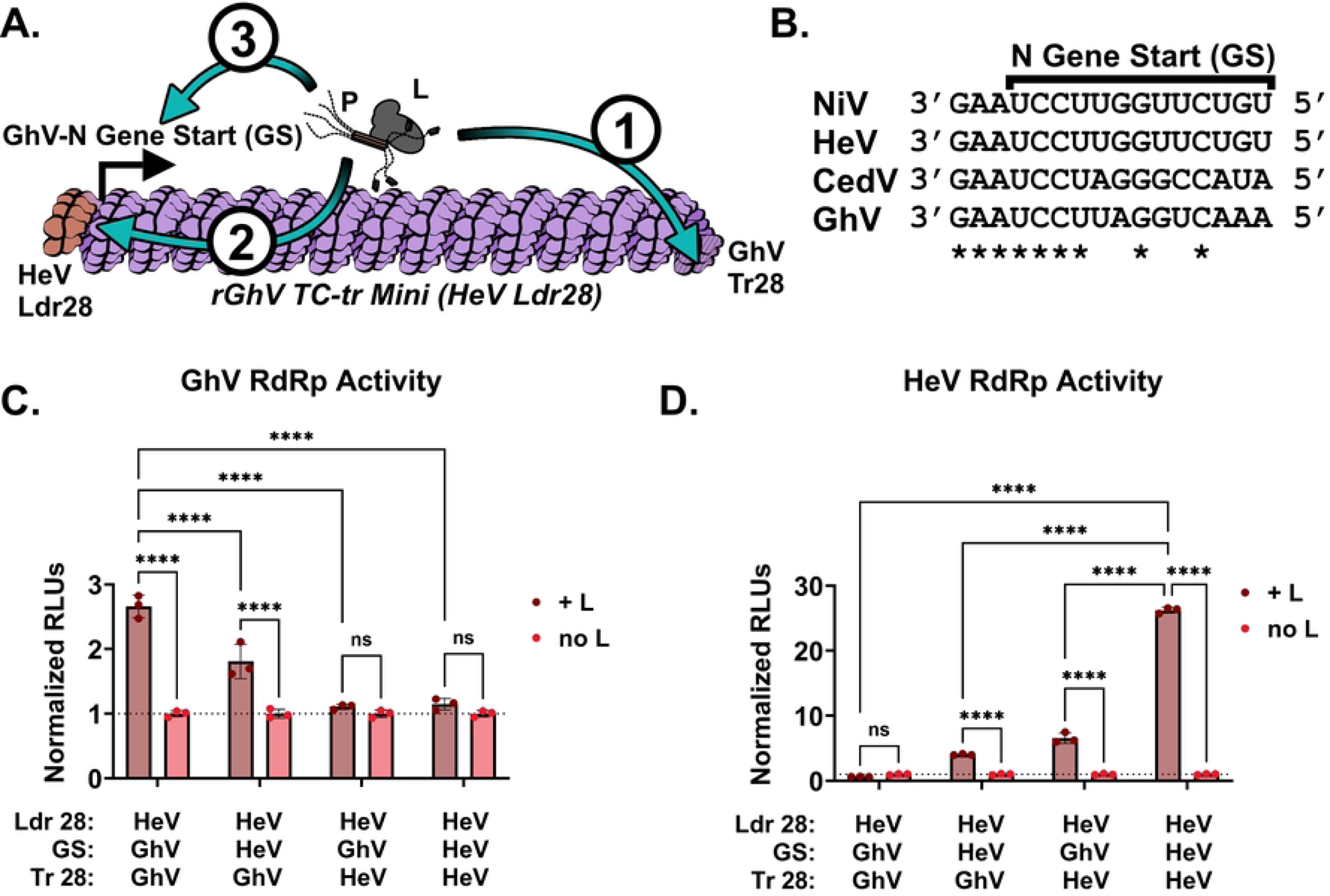
Incompatibilities exist between the respective replicase, gene starts, and 5’ Tr sequences of GhV and HeV. **(A**) Cartoon depicting the rGhV (HeV Ldr28) TC-tr minigenome and the order in which elements required for rescued must be recognized by the vRdRp. (1) The antigenomic promoter in the 5’ Tr must be recognized by the vRdRp to drive synthesis of genomic vRNA; (2) the genomic promoter in the 3’ Ldr of genomic vRNA must be recognized and must prompt the vRdRp to enter scanning mode; and (3) the N gene start must be recognized by the vRdRp to initiate transcription of viral mRNAs. **(B**) Alignment of the N gene start (GS) sequences from NiV, HeV, CedV, and GhV. Alignments are shown in the genomic vRNA sense, 3’ to 5’. Asterisks depict complete conservation. **(C)** GhV vRdRp activity above background resulting from the rescue of chimeric rGhV TC-tr minigenomes with systematic replacement of the GhV-N GS or Tr28 sequences with analogous HeV sequences. **(D)** HeV vRdRp activity above background resulting from the rescue of each chimeric rGhV TC-tr minigenome. RLUs from each rescue condition were normalized to RLUs derived from the rescue of respective minigenomes in the absence of HNV-L. Statistical significance was determined by a 2way ANOVA analysis in GraphPad Prism comparing normalized RLUs from rescues in the presence of HNV-L with normalized RLUs from rescues in the absence of HNV-L. Additional comparisons were conducted to determine significance between the parental rGhV (HeV Ldr28) TC-tr minigenome with the other minigenome mutants rescued in the presence of HNV-L. All rescues were conducted in triplicate. Error bars depict standard deviation. For all graphs (ns) = P > 0.05; (*) = P ≤ 0.05; (**) = P ≤ 0.01; (***) = P ≤ 0.001; (****) = P ≤ 0.0001.

### The vRdRp of Ghana virus is susceptible to two broad-spectrum antiviral compounds

Prior to the present study, the susceptibility of the GhV replicase to antivirals could not have been assessed, as infectious WT virus has not been isolated and the available genomic sequence was missing 28 nt from its 3’ Ldr. With the establishment of our GhV TC-tr minigenome system, we sought to test the susceptibility of the GhV replicase to EIDD-2749, a broad-spectrum vRdRp inhibitor, and GHP-88309, an allosteric inhibitor with efficacy against some paramyxovirus vRdRp(32, 33). To determine if TC-tr minigenomes could be reliably utilized to interrogate susceptibility of HNVs to antivirals, rNiV TC-tr VLPs were used to infect replicase-expressing BSRT7 cells in the presence of EIDD-2749. Nanoluciferase signal was inhibited by EIDD-2749 in rNiV TC-tr VLP infected cells, with an IC_50_ of 3.6 micromolar; inhibition was comparable to data collected using authentic rNiV at BSL-4, which yielded an IC_50_ of 2.6 micromolar. We likewise constructed and rescued a full-length rCedV clone encoding an eGFP reporter gene to compare inhibition of authentic virus with our rCedV TC-tr minigenome. Inhibition of the rCedV TC-tr minigenome with EIDD-2749 likewise recapitulated full-length virus susceptibility, with live virus demonstrating an IC_50_ of 0.7 micromolar and the rCedV TC-tr minigenome demonstrating an IC_50_ of 2.1 micromolar. Confident that our systems faithfully recapitulate authentic virus susceptibility to antiviral compounds, the GhV TC-tr minigenome was employed and demonstrated inhibition by EIDD-2749 with an IC_50_ of 1.0 micromolar (**Fig. 7A**).

**Figure 7.**
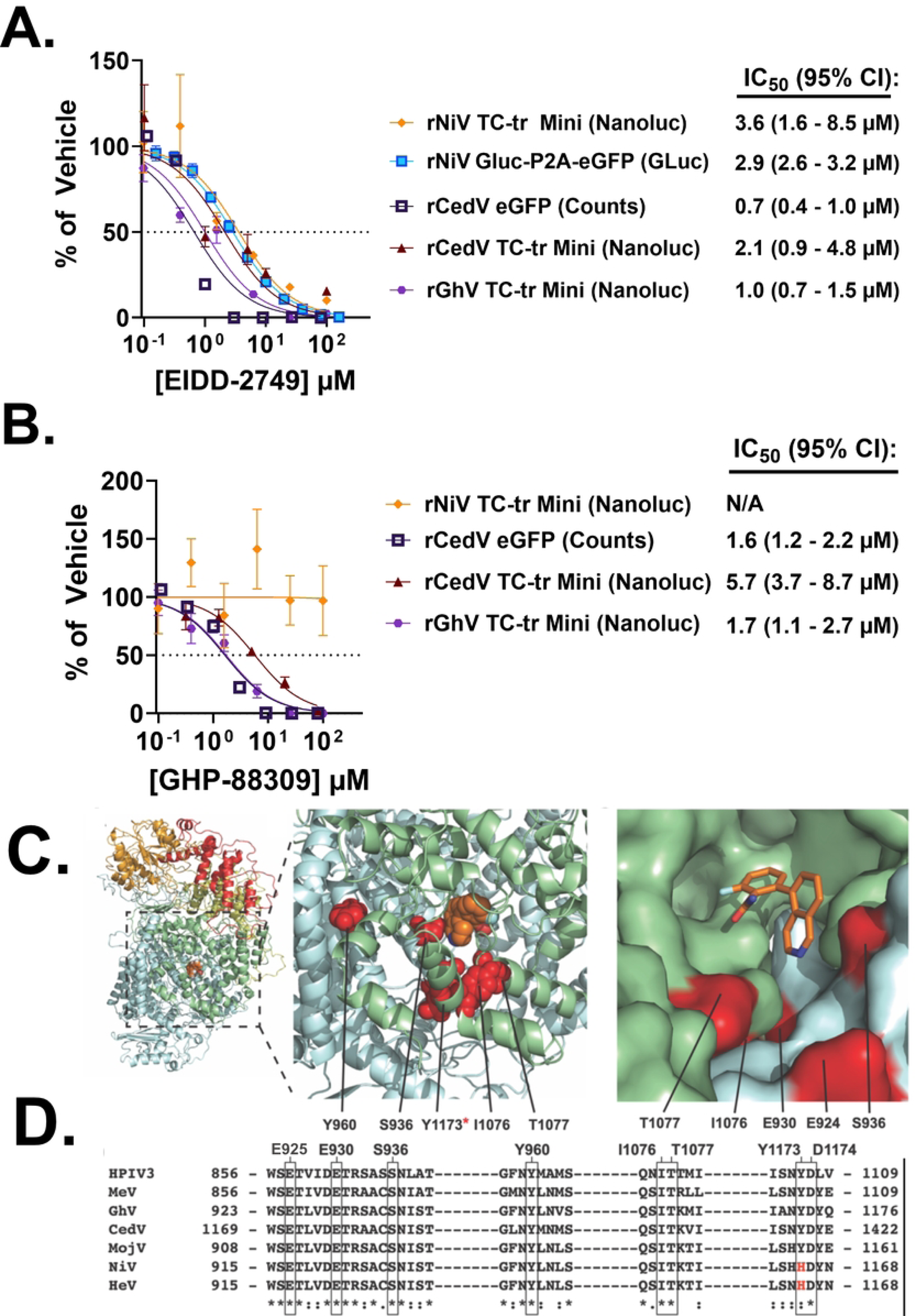
The GhV-L protein is susceptible to two vRdRp inhibitors. Inhibition curves conducted on a panel of authentic viruses or TC-tr minigenome systems treated with **(A)** EIDD-2749 or **(B)** GHP-88309. IC_50_ values were determined in GraphPad Prism by nonlinear regression of [inhibitor] vs. normalized response, and are listed to the right of each system implemented. The 95% confidence interval estimate for each IC_50_ is shown in parentheses. Error bars depict standard error of the mean. All inhibition experiments were conducted in at least biological triplicate. **(C)** Overview of a GhV-L homology model generated using PIV5-L as template (pdb: 6v85).The vRdRP, capping, connector, MTase, and CTD domains are colored blue, green, yellow, orange, and red, respectively. GHP-88309 is shown as orange spheres. The magnified inset depicts the locations where analogous residue mutations are known to induce resistance to GHP-88309. The third inset depicts alignment of a GHP-88309-MeV L complex with the GhV-L model (rms=0.373). Homologous residues to resistance sites in HPIV3 L are shown in red and labeled. GHP-88309 is shown as orange sticks. **(D)** Sequence alignment of HPIV3, MeV, and members of the HNV genus. All known residues shown to induce resistance (boxed) to GHP-88309 are conserved across genera except for NiV-L and HeV-L at position H1165 (red; H1165).

Inhibition assays utilizing GHP-88309 similarly revealed that the CedV vRdRp is potently inhibited, with IC_50_ values of 1.6 micromolar for authentic virus and 5.7 micromolar for the rCedV TC-tr minigenome. In agreement with previous studies, the rNiV TC-tr minigenome demonstrated no susceptibility to GHP-88309(33). GHP-88309 demonstrated a clear inhibition of GhV-L, with an IC_50_ of 1.7 micromolar (**Fig. 7B**). Homology modeling of GhV-L revealed a similar organization in the GhV vRdRp with other paramyxoviruses, and mapping of vRdRp mutations previously established to induce resistance to GHP-88309 onto this homology model suggests compatibility of GhV-L with GHP-88309 allosteric binding (**Fig. 7C**). In agreement with experimental results, sequence alignment of the L proteins from GHP-88309 susceptible species (HPIV3, MeV, CedV, and GhV) with resistant species (NiV and HeV) demonstrated that all residues previously shown to induce resistance to GHP-88309 are conserved across all aligned species except for NiV-L and HeV-L at position H1165 (**Fig. 7D**).

## Discussion

Noting the versatility of multicistronic minigenomes established for filoviruses, we sought to develop a comparable life-cycle modeling system for research on HNVs at BSL-2. Our efforts allowed us to successfully develop and rescue transcription and replication-competent (TC-tr) minigenomes for NiV, HeV, and CedV, demonstrating that such systems are broadly applicable to diverse HNVs (**Fig. 1**). TC-tr minigenomes provide a platform which enables a detailed examination of the molecular mechanisms underlying HNV replication and gene expression, including functions of the -M, -F, and - RBP genes in the context of viral genome replication and transcription.

Utilizing BiFC in the context of rNiV TC-tr minigenome infection enabled us to directly visualize the interaction of NiV-M, expressed under viral promoter, with host box C/D snoRNPs within syncytia. In agreement with previous reports that HNV-M localizes to nucleoli during infection, BiFC was observed in nucleolar punctae (**Fig. 2B, Supplementary Fig. 3F and 3G**). Because nucleolar trafficking of HNV-M is essential for budding, our BiFC observations in combination with detection of TC-tr VLPs in supernatant (**Fig. 2C**) demonstrates that NiV-M encoded by TC-tr minigenomes is functionally competent. The generation of syncytia driven by TC-tr minigenomes (**Fig. 1C and Fig. 2B**) likewise demonstrates that the encoded HNV-F and -RBP proteins are functional. In combination, HNV-M, -F, and -RBP under viral promoters could generate TC-tr VLPs that were competent for entry and could drive secondary infection in BSRT7 cells pre-transfected with the viral replicase (**Fig. 2D**). The observed reduction in titer for rNiV and rHeV TC-tr VLPs with iterative passaging may be due to delivering the replicase *in trans* via co-transfection, as not every cell will receive the optimal HNV-N, -P, and -L ratios. This limits the pool of cells capable of supporting robust minigenome activity and subsequent TC-tr VLP production. Future endeavors will seek to establish cell lines stably expressing one or more components of the viral replicase for each HNV.

The dearth of viral isolates for newly identified HNVs has severely limited our understanding of emerging species and their potential impacts on human health. Given the challenges posed by the absence of viral isolates, we sought to establish a TC-tr minigenome for GhV, a species which was identified solely through metagenomics(1). By constraining sequence alignments of the 3’ Ldr and 5’ Tr sequences to facilitate proper phasing of the bipartite promoter elements, we were able to determine that the reported sequence of GhV is missing 28 nucleotides from its 3’ Ldr sequence (**Fig. 3A**), but that its 5’ Tr sequence is complete (**Fig. 3B**). Proper phasing of the HNV bipartite promoters revealed a conservation of the core PrE-II sequence, consisting of the motif ‘ACC’ localized between hexamers 13 and 14, 14 and 15, and 15 and 16, respectively, which was previously characterized for NiV(22). The conservation of HNV PrE-II sequences lends insight as to why rescue of rHNV TC-tr minigenomes by heterotypic HNV replicases is possible, so long as PrE-I sequences are compatible with the vRdRp (**Fig. 5**). The unmapped sequence of GhV spans the first 5 hexamers, and thus GhV is effectively lacking a genomic PrE-I sequence (**Fig. 3A and 3D**). This poses a challenge for reverse genetics systems, as the GhV vRdRp will be unable to rescue a GhV minigenome based solely on its reported sequence.

To overcome the incomplete sequence of GhV, we replaced the unmapped region of the GhV 3’ Ldr with analogous sequences from NiV, HeV, or CedV (**Fig. 4A**). Implementing the NiV or HeV (but not CedV) Ldr28 sequences in lieu of the unmapped region of the GhV 3’ Ldr facilitated rescue of a GhV TC-tr minigenome by the GhV replicase **(Fig. 4B and 4C**). This work represents, to our knowledge, the first demonstration that the replicative machinery of GhV is functional. We observe that HNV antigenomic PrE-I sequences, when employed in lieu of genomic PrE-I sequences, lead to a marked reduction in reporter gene levels, indicating a loss in transcriptional activity by the vRdRp (**Fig. 4B-C, 4E-F**). This suggests that antigenomic PrE-I sequences may promote replicase activity of the vRdRp but are limited in prompting the vRdRp to enter scanning mode necessary for driving transcription. There are only 8 differences between the HeV Ldr28 and Tr28 sequences, with the first two hexamers being absolutely conserved (**Supplementary Fig. 7**). Because the first 12 nucleotides of the NiV 3’ Ldr have been demonstrated as sufficient for vRdRp engagement in a primer extension assay, our data suggests that sequences in hexamers 3-5 may be instrumental in influencing the triggering of the vRdRp into scanning mode(34). This aligns with existing models of paramyxovirus replication and transcription, where the viral antigenomic promoter would not be required to drive transcription, as no genes are encoded in the antigenomic vRNA(23, 24). While genomic and antigenomic promoters have been demonstrated to differentially modulate vRdRp transcriptional activity for other paramyxovirus species, such as Sendai virus, our findings extend these phenomena to the HNVs(35).

Heterotypic cross-rescue experiments can provide valuable insights into the genetic and functional relatedness between viral species(17, 36–38). We found that, as previously reported, the NiV and HeV replicases efficiently support rescue of each other’s minigenomes. However, neither replicase could support rescue of the rCedV nor rGhV TC-tr minigenomes (**Fig. 5**). This agrees with phylogenies demonstrating that NiV and HeV share a recent common ancestor relative to the other HNVs (**Supplementary Fig. 6**). Likewise, while the GhV replicase could recognize the genomic PrE-I sequences of NiV and HeV in the context of hybrid rGhV TC-tr minigenomes (**Fig. 4**), it was unable to support rescue of the WT rNiV, rHeV, or rCedV TC-tr minigenomes (**Fig. 5**). These findings suggest that as NiV, HeV, and GhV have diverged from their shared common ancestor, their respective vRdRps have likewise evolved distinct specificities for their respective templates.

Curiously, the CedV replicase demonstrated efficient recognition of the 3’ Ldr, GS, and 5’ Tr sequences of all HNVs tested, showcasing a broad plasticity in CedV vRdRp template recognition. The evolutionary trajectory of the CedV 3’ Ldr, GS, and 5’ Tr elements may have favored the selection of a vRdRp that requires a broader template recognition capacity in order to carry out its life cycle. Identifying the molecular determinants which confer the CedV vRdRp with its relative plasticity in template recognition will be the focus of future studies.

To determine at what level heterotypic cross-rescue is restricted between HeV and GhV, we constructed several rGhV (HeV Ldr28) TC-tr minigenomes in which we systematically replaced elements of the GhV sequence with their HeV equivalents (**Supplementary Fig. 5**). Rescue of these constructs revealed incompatibilities between the GhV repliase and the HeV antigenomic PrE-I (**Fig. 6C**); the HeV replicase could recognize the GhV-N GS sequence and the GhV antigenomic PrE-I sequences, albeit inefficiently, as changing either element individually supported rescue with a combinatorial positive effect (**Fig. 6D**). While recognition of individual, respective viral elements by a given vRdRp may not be completely incompatible, multiple inefficient compatibilities in combination can drive a total restriction of rescue. These findings offer insights on how GhV and HeV have diverged as genetic elements like antigenomic PrE-I or GS sequences change over evolutionary time, and that the vRdRp must likewise evolve to accommodate these changes.

We sought to leverage our TC-tr minigenomes to assess the susceptibility of emergent HNVs to orally efficacious broad-spectrum antivirals targeting the vRdRp. These include EIDD-2749, a nucleoside analogue effective against a wide array of RNA viruses, and GHP-88309, an allosteric inhibitor of the vRdRp with efficacy against a range of paramyxoviruses(32, 33). The GhV replicase was found to be potently inhibited by both EIDD-2749 and GHP-88309, with IC_50_ values of 1.0 micromolar and 1.7 micromolar, respectively (**Fig. 7A and 7B**). Authentic HNVs (rNiV and rCedV) and their TC-tr minigenome counterparts exhibited similar inhibition curves, demonstrating that antiviral susceptibility is comparably captured by TC-tr minigenomes. Thus, it is likely that authentic, full-length GhV should likewise be susceptible to both EIDD-2749 and GHP-88309. Homology modeling of the GhV-L protein in combination with sequence alignment of residues previously demonstrated to confer resistance against GHP-88309 provides a rational explanation for why GhV-L and CedV-L, but not NiV-L, are susceptible to GHP-88309 (**Fig. 7C and 7D**). Indeed, the H1165 residue in NiV-L was previously demonstrated in functional assay to confer resistance to GHP-88309(33). Because all susceptible paramyxovirus species have conserved residues at positions analogous to GhV-L E925, E930, S936, Y960, I1076, T1077, Y1173 (H1165 in NiV-L), and D1174, we propose that this association provides a means of predicting if a given paramyxovirus vRdRp will be susceptible to GHP-88309. Importantly, GHP-88309 has demonstrated a high oral bioavailability(33). This is valuable for pandemic preparedness and response efforts towards emergent paramyxoviruses.

While this study provides valuable insights into HNV biology, we acknowledge certain limitations that may influence the interpretation and generalizability of our findings. We acknowledge that the GhV vRdRp demonstrates relatively low activity above background as compared with the vRdRp of the other HNVs, and that this reduces the dynamic range of assays employing the GhV replicase. While all minigenome rescues were conducted with the same plasmid ratios/concentrations, relative expression profiles of individual HNV-N, -P, and -L were not assessed. The relatively low activity of the GhV replicase may be attributed to low expression of one or more components (GhV-N, -P, and/or -L), or could simply be reflective of kinetics specific to GhV. Further, we are aware that use of the HeV Ldr28 sequence may not be reflective of the authentic, unmapped 3’ Ldr sequence of GhV. While the HeV Ldr28 sequence provides a functional PrE-I which allows for GhV replicase activity, it may not be biologically identical to the unknown, authentic sequence.

In conclusion, this work showcases the value of TC-tr minigenomes as versatile, biologically contained life-cycle modeling systems for highly pathogenic HNVs. These systems are broadly applicable, allowing us to rescue diverse TC-tr minigenomes for NiV, HeV, and CedV. Further, application of this technology to GhV, a HNV which has never been isolated in culture, proved to be successful after restoration of a functional PrE-I sequence in its 3’ Ldr. The present study provides valuable insights into the replicative machinery and promoter recognition of GhV and lays the foundation for future efforts to rescue recombinant, full-length GhV with reverse genetics approaches. We further demonstrate that promoter recognition and GS recognition are distinct, yet combinatorially important processes that can dictate the successful rescue of viral species. Importantly, we have identified two vRdRp inhibitors which showcase potent inhibition of the GhV replicase, and a rationale for predicting if a given paramyxovirus polymerase is susceptible to GHP-88309. We are confident that TC-tr minigenomes and insights drawn from their implementation will undoubtedly bolster research and aid in pandemic preparedness and response efforts.

## Materials and methods

### Maintenance of Cell Lines

BSRT7/5, HeLa, BHK, and Vero CCL81 cells were maintained in Dulbecco’s modified Eagle medium (DMEM). Cells were grown at 37°C in 5% CO_2_. All media were supplemented with 10% fetal bovine serum which had undergone heat-inactivation at 56°C for 30 minutes.

### Design and cloning of reverse genetics plasmids

Fragments encoding the antigenome of rCedV with an eGFP reporter gene were synthesized (Twist Bioscience and GeneArt) and sequentially assembled into the pEMC vector. Briefly, the full-length construct was assembled by use of overlapping PCRs to join fragments using CloneAmp HiFi PCR mix (Takara Bio), with subsequent sequential insertions of joined fragments into restriction-digested pEMC vector by In-Fusion cloning (Takara Bio). The eGFP reporter gene is located between the N and P genes, and its insertion is accommodated by a duplication of the N-P intergenic sequences. The sequence of rCedV reflects strain CG1a (GenBank JQ001776.1). Recombinant HNV TC-tr minigenomes were designed as described in **Supplementary Fig. 1** and were assembled into a pcDNA3.1(−) vector lacking a CMV promoter, which was previously employed for our full-length rNiV reverse genetics system(39). For the rCedV TC-tr minigenome, primers were used to subclone respective elements of the viral sequence (ex: the hammerhead ribozyme and 3’ Ldr sequence, the sequences between start codon of CedV-M and the stop codon of CedV-RBP, and the 5’ Tr sequence and HDV ribozyme) from the full-length rCedV antigenomic plasmid. Previously established rNiV and rHeV reverse genetics plasmids were used as templates for subcloning their respective elements, as well as for deriving the mCherry reporter gene. The rNiV TC-tr minigenome reflects strain UMMC1 (GenBank AY029767.1) and the rHeV TC-tr minigenome reflects strain HeV/Australia/1994/Horse18 (MN062017.1) The full antigenomic sequence of rGhV strain Eid_hel/GH-M74a/GHA/2009 (NCBI Reference Sequence NC_025256.1) was fully synthesized (Bio Basic Inc.) and was used as template for subcloning the respective GhV sequences. All respective HNV fragments were assembled by use of overlapping PCRs to join fragments using CloneAmp HiFi PCR mix (Takara Bio), with subsequent assembly into the MluI/PmeI digested pcDNA3.1(−) vector by In-Fusion cloning (Takara Bio). The viral antigenomic sequences in each reverse genetics construct is flanked on its 5’ terminal end by a sequence-specific hammerhead ribozyme element and on its 3’ terminal end by the HDV ribozyme, as has been detailed for our paramyxovirus reverse genetics systems(25, 26, 39). An optimal T7 promoter sequence lies upstream of the 5’ hammerhead ribozyme sequence, and a T7 terminator sequence lies downstream of the 3’ HDV ribozyme sequence. The accessory plasmids for NiV (NiV-N, -P, and -L) and HeV (HeV-N, -P, and -L) were constructed and described previously(39). Accessory plasmids encoding CedV-N, -P, and -L, or GhV-N, -P, and -L, respectively, were subcloned from their respective viral antigenome plasmids and incorporated into the pTM1 vector between the SpeI and XhoI restriction sites, as previously described for NiV and HeV accessory plasmid construction(17). All restriction enzymes were purchased from New England Biolabs, and all primers employed were synthesized by Millipore Sigma.

### Rescue of rHNV TC-tr minigenomes

BSRT7/5 cells were seeded one day prior to transfection in 12-well or 24-well format, to achieve 70-80% confluence on the day of transfection. Twenty-four hours later, cells were co-transfected with plasmids encoding respective the rHNV TC-tr minigenome, codon-optimized T7 polymerase, and HNV-N, -P, and -L as previously described for full length HNV rescue, but adapted for smaller well format(25). All rescue transfections used Lipofectamine LTX and PLUS according to the manufacturer recommendations. For 12-well format this equated to 2.55 micrograms of rHNV TC-tr minigenome, 0.73 micrograms of codon-optimized T7 polymerase, 0.91 micrograms of HNV-N, 0.58 micrograms of HNV-P, 0.29 micrograms of HNV-L, 2.33 microliters of PLUS, and 3.76 microliters of LTX per reaction. For 24-well format this equated to 1.39 micrograms of rHNV TC-tr minigenome, 0.40 micrograms of codon-optimized T7 polymerase, 0.49 micrograms of HNV-N, 0.32 micrograms of HNV-P, 0.16 micrograms of HNV-L, 1.27 microliters of PLUS, and 2.04 microliters of LTX per reaction. For all rescue transfections, mastermixes were prepared which contained the TC-tr minigenome, HNV-N, HNV-P, and codon-optimized T7; these master mixes were then aliquoted into equal parts before adding plasmid encoding HNV-L or plasmid encoding GFP. Plasmid encoding GFP was used in lieu of HNV-L as a control to determine background signal nonspecific to the viral replicase. Twenty-four HPT, media was replaced on plates. Quantification of rescue events and nanoluciferase assays were conducted at 48-72 hours post-transfection, depending on the experiment.

### Production and propagation of TC-tr VLPs

Rescue was performed as described, with a media change 24 hours post transfection. TC-tr VLP containing supernatant was then harvested at 72 hours post transfection for use in downstream experiments. Aliquots of TC-tr VLPs were stored at −80°C. BSRT7/5 cells were seeded to 70-80% confluence in 24-well format and co-transfected with 0.50 micrograms of HNV-N, 0.32 micrograms of HNV-P, 0.16 micrograms of HNV-L, and 0.40 micrograms of codon-optimized T7 polymerase using BioT reagent (Bioland) according to manufacturer instructions. Twenty-four HPT, 125 microliters of undiluted TC-tr VLP containing supernatant was used to infect the cells in low volume for one hour at 37°C, with gentle rocking every 15 minutes. Media was then brought to 0.5 mL per well. Cells were monitored for mCherry events and imaged at 72 hours post infection. Supernatant was harvested from passage 1 (P1) cells at 72 hours post infection and used to infect BSRT7/5 cells in identical manner for passage 2 (P2). For infection of BSRT7/5 cells with HiBiT-mCherry constructs, two wells of a six well plate were seeded with four-hundred thousand BSRT7/5 cells. After twenty-four hours, cells were co-transfected with 0.72 micrograms of HNV-N, 0.46 micrograms of HNV-P, 0.23 micrograms of HNV-L, and 0.58 micrograms of codon-optimized T7 polymerase using BioT reagent (Bioland). Six hours post-transfection, 500 microliters of undiluted HiBiT-mCherry TC-tr VLP-containing rescue supernatant was used to infect the six well plates via spinoculation at 2,000 RPM for 1 hour at 37°C. Following spinoculation, cells were given a 1 hour recovery at 37°C prior to resuspension by trypsin-EDTA (0.25%). Cells were seeded equally into 96-well format directly into drug-containing media for inhibition assays.

### Microscopy and quantification of rHNV TC-tr minigenome mCherry events

At 72 hours post-transfection, TC-tr minigenome rescue cells were stained with Hoechst (Abcam ab228551) for approximately 10 minutes prior to a complete media change. Images of TC-tr minigenome rescue events were then immediately captured on a Cytation 3 plate reader (BioTek) using the blue and red channels. All images were exported to ImageJ for processing. Images shown in the same figure panel were captured in the same experiment and were processed with identical conditions. A Celigo Imaging Cytometer (Nexcelom) was used to image the entire plate of TC-tr minigenome rescues in the red channel. The total number of mCherry positive events were quantified using Celigo software analysis of the red channel.

### Quantification of vRdRp activity above background

To measure nanoluciferase activity, TC-tr minigenome rescue cells were lysed and processed using the Nano-Glo® HiBiT Lytic Detection System (Promega Corporation), according to the manufacturer recommendations. Cell lysate was transferred to white 96-well plates and RLUs were measured on a Cytation 3 plate reader (BioTek) using a gain of 125 and an integration time of 1.0s. Raw RLUs were then normalized to the average RLUs of the no-L (GFP) control to determine the vRdRp-dependent signal above background.

### Cloning of BiFC constructs

Total RNA was extracted from HEK-293 cells using a Direct-zol RNA miniprep kit (Zymogen) and was used as template for RT-PCR to derive cDNA of human NOP56, NHP2L1, and FBL. One microgram of RNA was used as template for the reactions conducted using the SuperScript™ III One-Step RT-PCR System with Platinum™ Taq DNA Polymerase (Invitrogen). Gene-specific primer sequences were employed for deriving cDNA of the CDS of NOP56 (primer FWD 5’- ATGGTGCTGTTGCACGTG-3’ and primer REV 5’- CTAATCTTCCTGGGATGCTTTATG-3’), NHP2L1 (primer FWD 5’- ATGACTGAGGCTGATGTGAA-3’ and primer REV 5’- TTAGACTAAGAGCCTTTCAATGGAC-3’) and FBL (primer FWD 5’- ATGAAGCCAGGATTCAGTCC-3’ and primer REV 5’- TCAGTTCTTCACCTTGGGG-3’). CDS cDNA was then used as template for PCR with HiFi PCR mix (Takara Bio) to make each ORF compatible for In-Fusion cloning (Takara Bio). Sequences were assembled into pcDNA3.1 vector encoding VC155 or VN173 on the N- or C-terminus, ultimately fused to the CDS of the host gene by a 3x GGGSG linker. BiFC constructs for NiV-M were previously described(29). The rNiV TC-tr minigenome encoding VC155 was constructed by subcloning the VC155 sequence from the pcDNA3.1 VC155-2xLinker-NiV-M construct and assembling it into restriction-digested rNiV TC-tr minigenome by In-Fusion cloning (Takara Bio) approaches. All primers were synthesized by Millipore Sigma.

### Bimolecular fluorescence complementation assay

One hundred thirty-five thousand HeLa or BSRT7/5 cells were seeded into a 12-well plate. Twenty-four hours later, BSRT7/5 cells were transfected with rNiV TC-tr minigenomes encoding WT (untagged) NiV-M, or NiV-M with VC155 fused to its N terminus. In parallel, HeLa cells were transfected with 1.5 micrograms of either NOP56 or NHP2L1 fused on their respective C termini with VN173. Twenty-four hours post-transfection, BSRT7/5 and HeLa cells were resuspended using trypsin-EDTA (0.25%), and equal amounts of BSRT7/5 and HeLa cells from each transfection condition were co-cultured in different combinations in six-well format. At 72 hours post-transfection, cells were stained with Hoechst (Abcam ab228551) for approximately 10 minutes prior to a complete media change. Images were captured on a Cytation 3 plate reader (BioTek) using the green, red, and blue channels. All images were exported to ImageJ for processing. Images shown in the same figure panel were captured in the same experiment and were processed with identical conditions.

### Detection of viral genomes in supernatant

Supernatant was harvested from TC-tr rescue cells and was clarified by brief centrifugation. One hundred microliters of supernatant was used for RNA extraction using the QIAamp® Viral RNA Mini kit (QIAGEN). Sample was treated with DNAse according to manufacturer recommendations, to remove any residual antigenome rescue plasmid from the system. RNA was eluted in ultra-pure water and stored at −80°C. The Primer-free LunaScript® RT Master Mix Kit (New England Biolabs) was used for first strand cDNA synthesis using a gene-specific primer targeting the mCherry gene in the genomic vRNA (primer sequence 5’-CGCTTCAAGGTGCACATGG-3’). Following cDNA synthesis, equal volumes of cDNA were used for qPCR using the Luna® Universal qPCR Master Mix kit (New England Biolabs). For qPCR, gene specific primers were employed targeting a 270 basepair region of the mCherry gene (sequences Primer FWD 5’-CGCTTCAAGGTGCACATGG-3’ and Primer REV 5’- GCCGTCCTCGAAGTTCATCAC-3’). The rHeV TC-tr minigenome plasmid was serially diluted and used as a standard to calculate cDNA copy number. Signal was captured on a C1000 Touch Thermal Cycler (Bio-Rad) and data was exported for analysis. All qPCR reactions were conducted in technical duplicate for each biological sample.

### Rescue of rCedV encoding eGFP

Four-hundred thousand BSRT7/5 cells were seeded into each well of a six well plate. Twenty-four hours later 7.0 micrograms of plasmid encoding the rCedV eGFP antigenome, 2.0 micrograms of plasmid encoding codon-optimized T7 polymerase, 2.5 micrograms of pTM1-T7-CedV-N, 1.6 micrograms of pTM1-T7-CedV-P, and 0.8 micrograms of pTM1-T7-CedV-L were diluted in 200 microliters of OptiMEM and gently mixed. Transfections were conducted using 6.4 microliters of Lipofectamine PLUS reagent and 10.3 microliters of Lipofectamine LTX reagent. After a 30 minute incubation at room temperature, transfection complexes were added dropwise to BSRT7/5 cells. Media was changed 24 hours post-transfection and cells were monitored daily for GFP events and secondary spread within the well. To amplify virus, two T175 flasks of 75% confluent Vero CCL81 cells were infected with 250 microliters of rescue supernatant. By 48 hours post-infection, the monolayer was completely infected, with widespread GFP signal and cytopathic effect. Supernatant was harvested, clarified by brief centrifugation, and frozen at −80°C. Virus was titrated on Vero CCL81 cells by serial dilution in 96-well format; 24 hours post-infection, the plate was imaged using a Celigo Imaging Cytometer (Nexcelom) in the green channel. The total number of GFP positive events were quantified using Celigo software analysis of the green channel.

### Antiviral compounds and inhibition assays

Both EIDD-2749 and GHP-88309 were resuspended to 100mM in DMSO and aliquoted in small volume to avoid multiple freeze-thaws. Compounds were stored at −80°C until use. Vehicle (DMSO) alone was used as a control for all assays. For TC-tr minigenome inhibition assays, BSRT7/5 cells were pre-transfected and infected with TC-tr VLPs as described, or BSRT7/5 cells were transfected for TC-tr minigenome rescue as described, and were trypsinized and equally seeded into media containing respective compound dilutions. Media was replaced at 24 hours post-treatment with drug-containing media or DMSO control to ensure compound integrity was not compromised before reporter gene readout. At 48 hours post-treatment, cells were lysed for nanoluciferase assay. Percent activity of vehicle was calculated for each condition by normalizing all values to the average of the DMSO control and calculating 100*(sample normalized RLUs). To account for the narrow dynamic range of the rGhV TC-tr minigenome, a no-L control was included to measure vRdRp-independent background, which was subtracted from the raw RLUs of each condition. Any negative values were set to zero, and the percent of vehicle was then calculated as described above using the background-subtracted RLUs.

For authentic rCedV experiments, 11,000 cells/well of BSRT7/5 cells were seeded in 96-well format, and infected the next day with rCedV eGFP at an MOI of 0.2. Compounds were added at the time of infection. At 24 hours post-infection, infected plates were captured using a Celigo Imaging Cytometer (Nexcelom) in the green channel. Celigo analysis software was then employed to quantify the number of events in the green channel of each well. The rNiV (Malaysia strain UMMC1) encoding eGFP and *Gaussia* luciferase (GLuc) was rescued as previously described(12, 39, 40). All experiments conducted with authentic rNiV were conducted in a class II BSC at the Galveston National Laboratory BSL-4 at the University of Texas Medical Branch (UTMB). For rNiV inhibition assays, 11,000 cells/well of BHK cells were seeded in 96-well format and infected the next day with rNiV encoding a GLuc-P2A-eGFP reporter at an MOI of 0.05. At 24 HPI, supernatant was collected and transferred to a white plate for GLuc assay using the Pierce™ *Gaussia* Luciferase Glow Assay Kit. A Cytation 5 plate reader (BioTek) was used to measure RLUs from viral supernatant. For all authentic virus experiments, raw RLUs or GFP counts were normalized to the DMSO control, and then percent of vehicle was calculated as described above. All drug inhibition assays were conducted in at least biological triplicate, and IC_50_ values were calculated in GraphPad Prism software by nonlinear regression of the [inhibitor] vs normalized response.

## Acknowledgements

This work was supported in part by NIH grants R01 AI071002 (B.L., R.K.P.), U19 AI171403 (B.L., R.K.P., R.C., A.N.F.), R21 AI149033 (B.L.). B.L. also acknowledges support from the Bill & Melinda Gates Foundation (PAD Henipavirus Initiative, INV-048877). K.N.J. was supported by a NIH T32 Biodefense training fellowship (AI060549). K.A. was supported by a NIH T32 training fellowship (AI07647). G.D.H. is supported by the National Science Foundation Graduate Research Fellowship Program (Grant No. 1842169). Any opinions, findings, and conclusions or recommendations expressed in this material are those of the author(s) and do not necessarily reflect the views of the National Science Foundation.

## Supporting Information

**Supplementary Figure 1.**
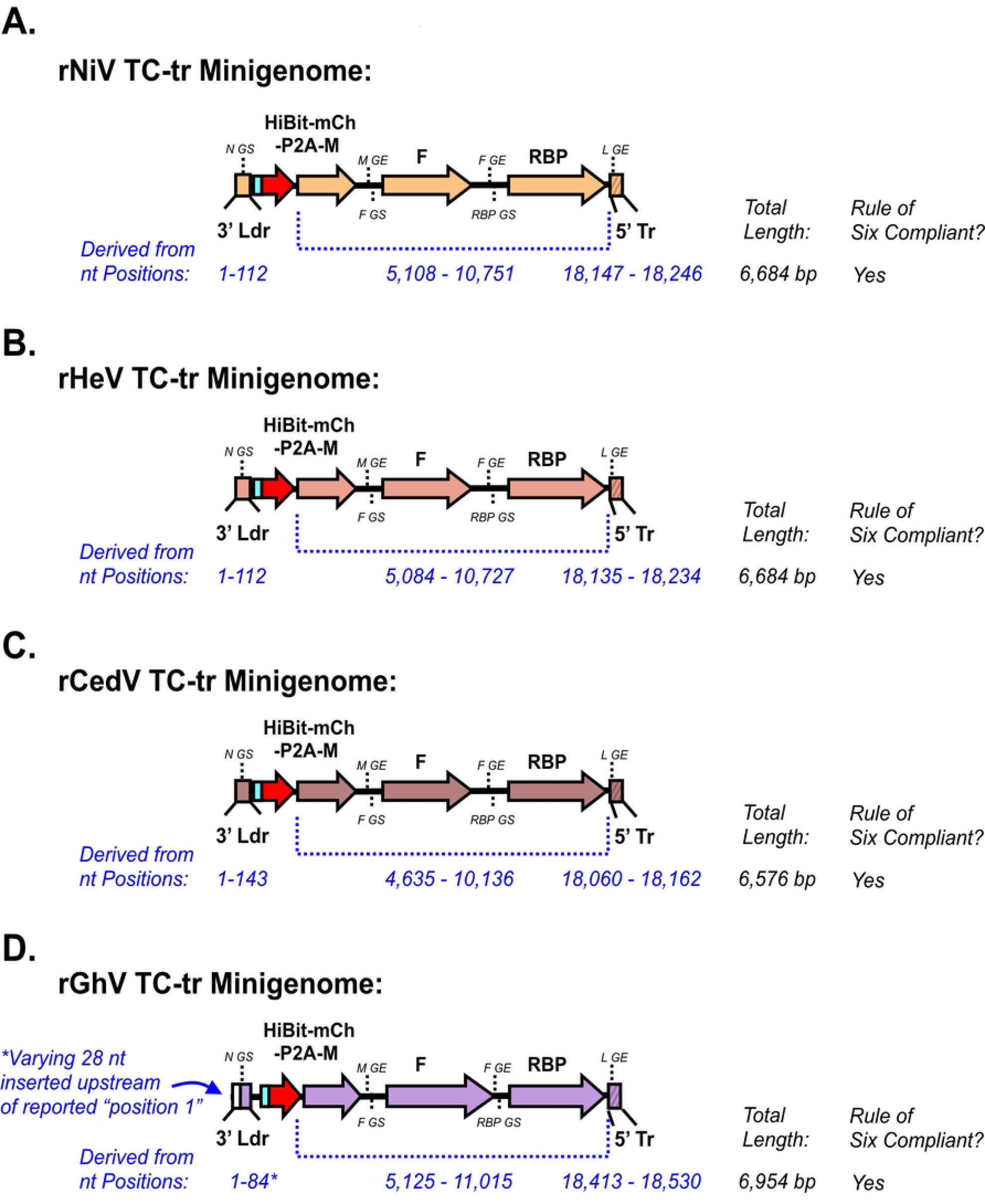
Schematic representation of rHNV TC-tr minigenome construct designs. Tetracistronic henipavirus minigenomes were designed for **(A)** Nipah virus, **(B)** Hendra virus, **(C)** Cedar virus, **(D)** and Ghana virus. All constructs encode the respective reported 3’ Ldr sequence through the start codon of the N gene, the viral sequence spanning the start codon of the M gene through the stop codon of the RBP gene, and at least 100 nucleotides derived from the 5’ Tr sequence. All constructs were designed to be compliant with the rule of six; if required, additional stop codons were added after the RBP gene to fulfill this rule. Each panel details which respective regions of the viral genome were assembled together, with nucleotide positions corresponding to the accession sequence: For NiV, derived sequences are from strain UMMC1 (GenBank AY029767.1); for HeV, derived sequences are from HeV/Australia/1994/Horse18 (MN062017.1); for CedV, derived sequences are from strain CG1a (GenBank JQ001776.1); and for GhV, derived sequences are from strain Eid_hel/GH-M74a/GHA/2009 (NCBI Reference Sequence NC_025256.1). Note that for GhV (**D**), an additional 28 nucleotides are inserted upstream of the reported 3’Ldr sequence for all constructs.

**Supplementary Figure 2.**
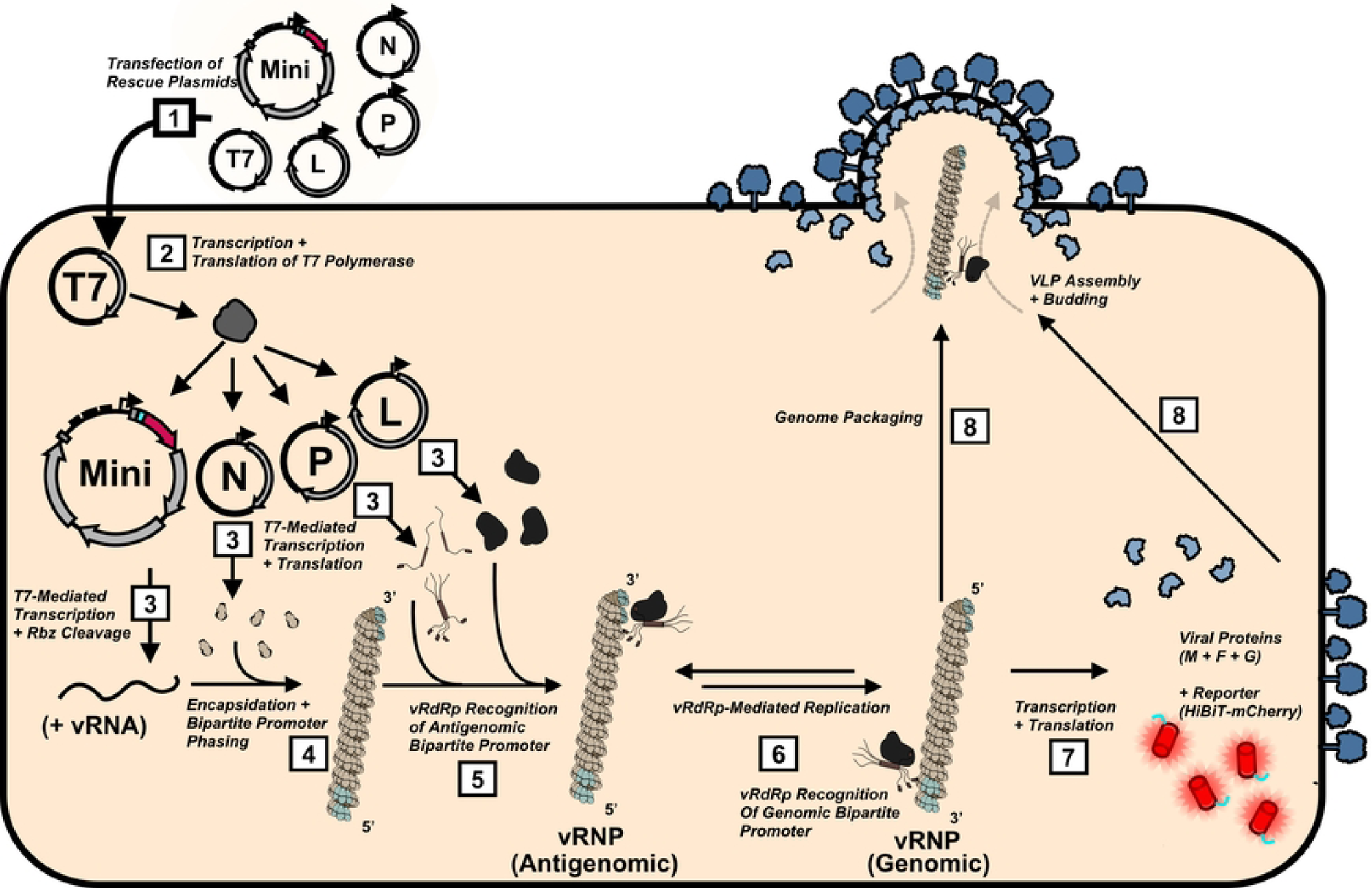
Schematic detailing TC-tr minigenome rescue. For minigenome rescue to occur, plasmids encoding the rHNV TC-tr minigenome, T7-HNV-N, T7-HNV-P, T7-HNV-L, and codon-optimized T7 polymerase must be cotransfected into cells (1). Following co-transfection, T7 polymerase is transcribed and translated (2). T7 polymerase drives transcription of HNV-N, -P, -L, and the rHNV TC-tr minigenome in the cellular cytoplasm. As the rHNV TC-tr minigenomic RNA is transcribed, the ribozyme elements at the terminal ends fold and cleave, resulting in antigenomic vRNA without exogenous sequence on the ends (3). HNV-N binds to and oligomerizes along the length of the minigenomic vRNA, resulting in the three-dimensional phasing of the antigenomic bipartite promoter onto the same surface of the vRNP (4). Upon proper phasing of the antigenomic bipartite promoter, HNV-P may recruit HNV-L to the vRNP, forming the replicase (5). HNV-L may then recognize the antigenomic bipartite promoter, resulting in vRdRp dependent replication and genomic minigenomic vRNA. Proper phasing of the genomic bipartite promoter results in recognition by the vRdRp, resulting in further rounds of replication (6) and transcription (7) of genes encoded by the minigenomic vRNA. Expression of HiBiT-mCherry provides a visual and quantitative readout for vRdRp activity, and expression of HNV-M, -F-, and -RBP can result in packaging of the minigenomic vRNP into TC-tr VLPs (8).

**Supplementary Figure 3.**
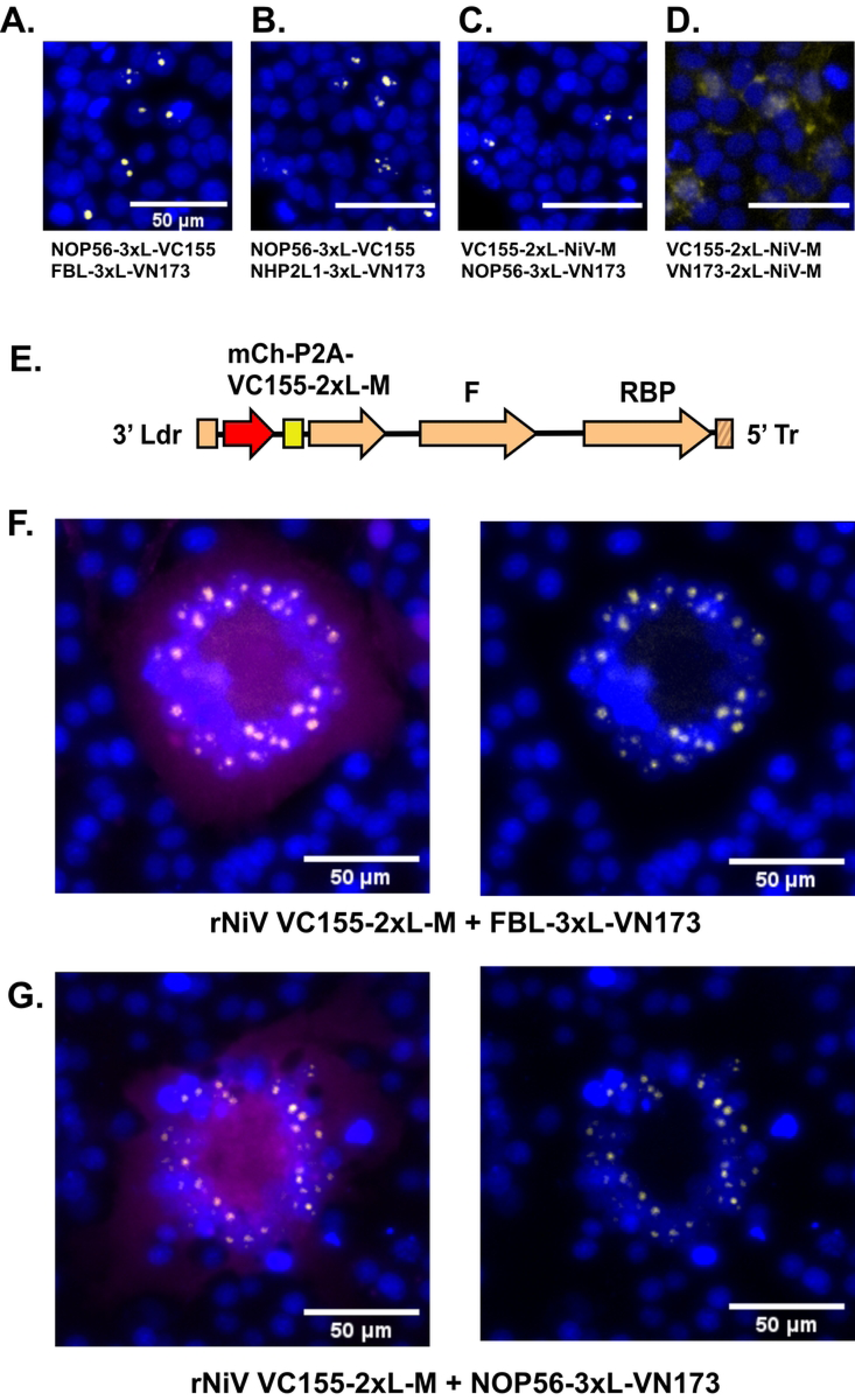
Validation of BiFC constructs. Co-transfection of HEK-293T cells with (A) NOP56-3xL-VC155 and FBL-3xL-VN173, or (B) NOP56-3xL-VC155 and NHP2L1-3xL-VN173 results in BiFC signal exclusively in nucleolar punctae. Further, co-transfection of HEK-293T cells with (C) VC155-2xL-NiV-M with NOP56-3xL-VN173, but not (D) VC155-2xL-NiV-M with VN173-2xL-NiV-M, yields BiFC in nucleoli. (E) Design of a rNiV TC-tr minigenome encoding VC155 tethered to the N-terminus of NiV-M, with the two separated by a 2x GGGGS linker. Rescue of the rNiV TC-tr minigenome encoding VC155-2xL-NiV-M in BSRT7 cells co-transfected with (F) FBL-3xL-VN173 or (G) NOP56-3xL-VN173 yields mCherry-positive syncytia with BiFC localized in nucleolar punctae. Both (F) and (G) show panels with mCherry signal (left) and without (right) for better visualization of nuclei and BiFC.

**Supplementary Figure 4.**
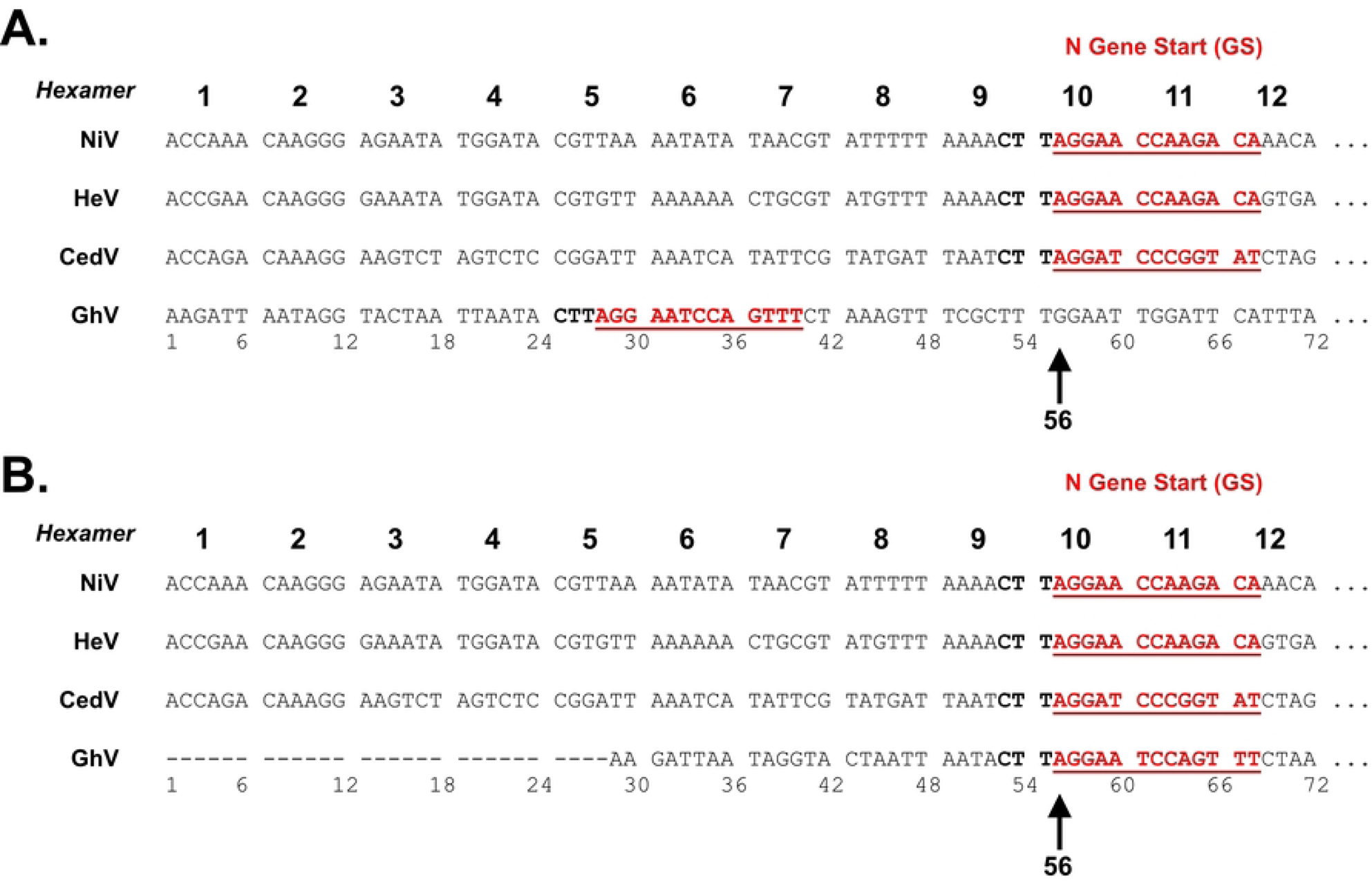
Accounting for unmapped nucleotides in the GhV 3’ Ldr sequence properly phases the GhV N gene start. (A) Uncorrected sequence alignment of the first 72 nucleotides of the 3’ Ldr sequences of NiV, HeV, and CedV with the reported 3’ Ldr sequence of GhV (all shown as cDNA, 5’ to 3’). (B) Shifting the reported GhV 3’ Ldr sequence by 28 nucleotides effectively positions its N gene start at position 56; hexamer phasing of the N gene start is invariantly conserved among the fully-sequenced bat-borne henipaviruses. For all alignments, the N gene start is colored in red.

**Supplementary Figure 5.**
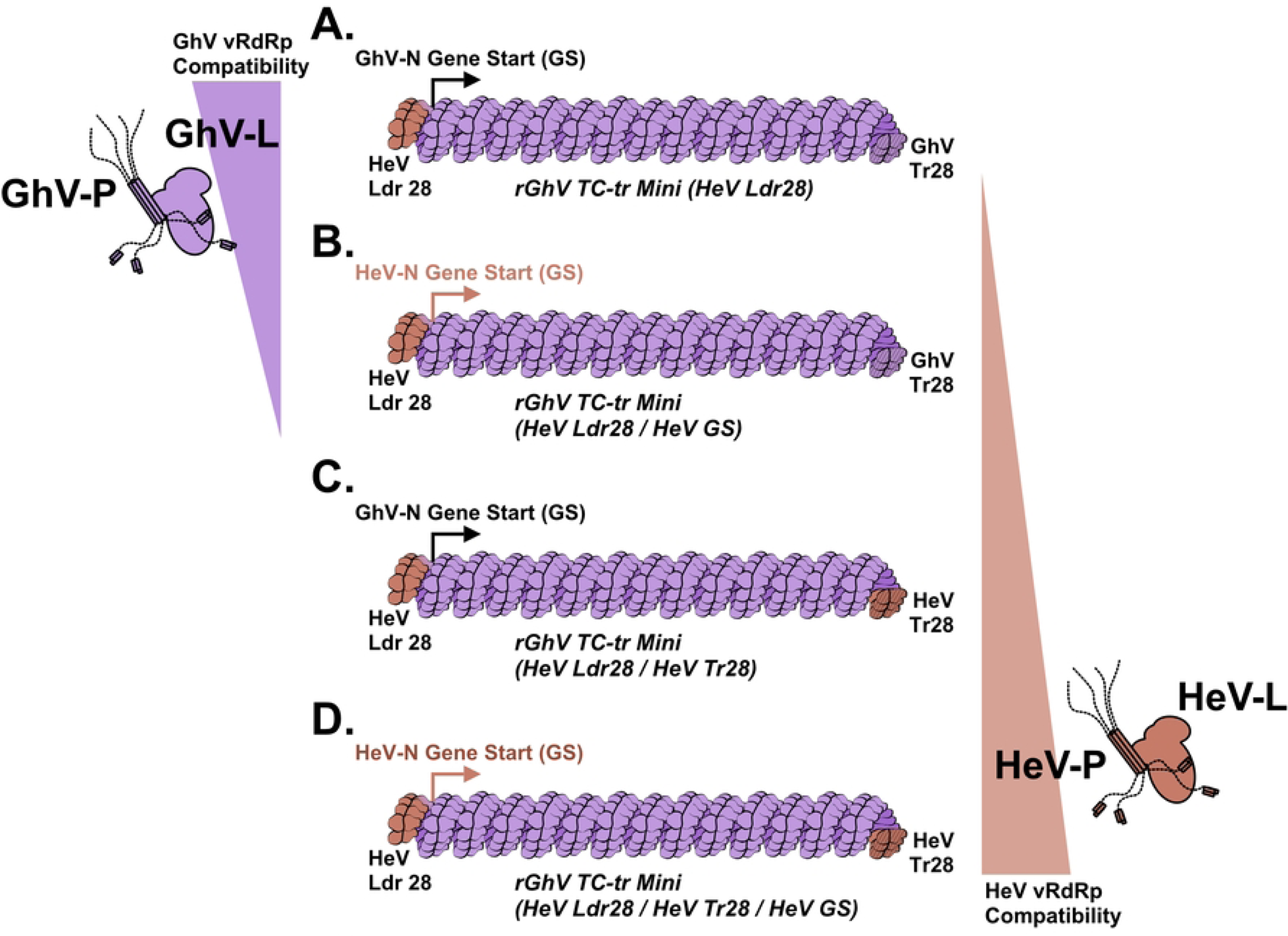
Depiction of rGhV (HeV Ldr28) TC-tr minigenomes with systematic replacement of the GhV-N GS or Tr28 nucleotides with HeV equivalents. Schematic demonstrates various minigenomes relative to (A) the parental rGhV TC-tr minigenome (HeV Ldr28) construct. (B) Replacement of either the GhV-N GS with the HeV-N GS, or (C) replacement of the terminal 28 nucleotides of the GhV 5’Tr (Tr28) with the HeV equivalent. (D) Combined replacement of both the GhV-N GS and GhV Tr28 sequences with the HeV equivalents. On the left, cartoon signifying which constructs support GhV replicase activity. On the right, carton signifying which constructs support HeV replicase activity.

**Supplementary Figure 6.**
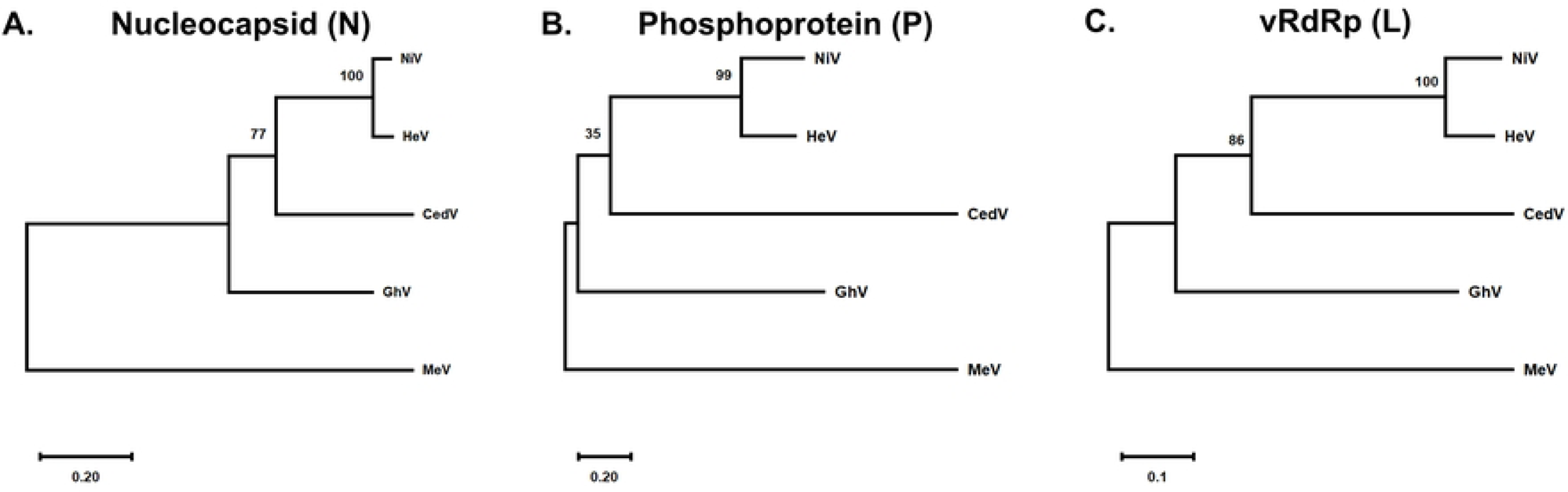
Phylogenetic relatedness of bat-borne HNV replicase proteins. Phylogenetic trees demonstrating relatedness between the NiV, HeV, CedV, and GhV (A) nucleocapsid, (B) phosphoprotein, and (C) large (vRdRP) proteins. For all analyses, measles virus strain IC323 (MeV) is used as an outgroup. The evolutionary history was inferred by using the Maximum Likelihood method and JTT matrix-based model. The tree with the highest log likelihood is shown. The percentage of trees in which the associated taxa clustered together is shown next to the branches. Initial tree(s) for the heuristic search were obtained automatically by applying Neighbor-Join and BioNJ algorithms to a matrix of pairwise distances estimated using the JTT model, and then selecting the topology with superior log likelihood value. The tree is drawn to scale, with branch lengths measured in the number of substitutions per site. These analyses involved 5 amino acid sequences. Evolutionary analyses were conducted in MEGA X.

**Supplementary figure 7.**
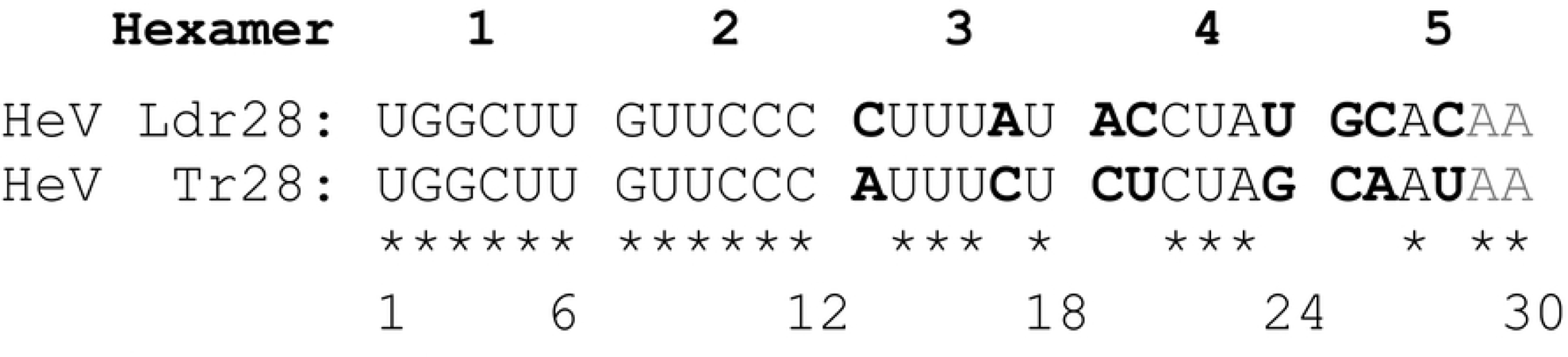
Sequence alignment of HeV Ldr28 and Tr28 sequences. Sequence alignment of the HeV Ldr28 and Tr28 sequences employed in this study. Differences between the two sequences are localized to nucleotide positions 13 and 17 in hexamer 3; positions 19, 20, and 24 in hexamer 4; and positions 25, 26, and 28 in hexamer 5.

## References

1. Drexler JF, Corman VM, Müller MA, Maganga GD, Vallo P, Binger T, et al. Bats host major mammalian paramyxoviruses. Nat Commun. 2012;3:796.

2. Barr J, Smith C, Smith I, De Jong C, Todd S, Melville D, et al. Isolation of multiple novel paramyxoviruses from pteropid bat urine. Journal of General Virology. 2015;96(1):24–9.

3. Larsen BB, Gryseels S, Otto HW, Worobey M. Evolution and Diversity of Bat and Rodent Paramyxoviruses from North America. Journal of Virology. 2021-10-20;96(3).

4. Wells HL, Loh E, Nava A, Solorio MR, Lee MH, Lee J, et al. Classification of new morbillivirus and jeilongvirus sequences from bats sampled in Brazil and Malaysia. Archives of Virology 2022 167:10. 2022-07-04;167(10).

5. Eaton BT, Broder CC, Middleton D, Wang L-F. Hendra and Nipah viruses: different and dangerous. Nat Rev Microbiol. 2006;4(1):23–35.

6. Hendra and Nipah viruses: why are they so deadly? Current Opinion in Virology. 2012/06/01;2(3).

7. Devnath P, Wajed S, Chandra Das R, Kar S, Islam I, Masud HMAA. The pathogenesis of Nipah virus: A review. Microbial Pathogenesis. 2022;170:105693.

8. Gurley ES, Montgomery JM, Hossain MJ, Bell M, Azad AK, Islam MR, et al. Person-to-Person Transmission of Nipah Virus in a Bangladeshi Community. Emerging Infectious Diseases. 2007/07;13(7).

9. Schountz T, Campbell C, Wagner K, Rovnak J, Martellaro C, Debuysscher B, et al. Differential Innate Immune Responses Elicited by Nipah Virus and Cedar Virus Correlate with Disparate In Vivo Pathogenesis in Hamsters. Viruses. 2019;11(3):291.

10. Lieu KG, Marsh GA, Wang L-F, Netter HJ. The non-pathogenic Henipavirus Cedar paramyxovirus phosphoprotein has a compromised ability to target STAT1 and STAT2. Antiviral Research. 2015;124:69–76.

11. Marsh GA, De Jong C, Barr JA, Tachedjian M, Smith C, Middleton D, et al. Cedar Virus: A Novel Henipavirus Isolated from Australian Bats. PLoS Pathogens. 2012;8(8):e1002836.

12. Pernet O, Schneider BS, Beaty SM, LeBreton M, Yun TE, Park A, et al. Evidence for henipavirus spillover into human populations in Africa. Nature Communications. 2014;5(1):5342.

13. Madera S, Kistler A, Ranaivoson HC, Ahyong V, Andrianiaina A, Andry S, et al. Discovery and Genomic Characterization of a Novel Henipavirus, Angavokely Virus, from Fruit Bats in Madagascar. Journal of Virology. 2022/09;96(18).

14. Chua KB, Lek Koh C, Hooi PS, Wee KF, Khong JH, Chua BH, et al. Isolation of Nipah virus from Malaysian Island flying-foxes. Microbes and Infection. 2002;4(2):145–51.

15. Halpin K, Young PL, Field HE, Mackenzie JS. Isolation of Hendra virus from pteropid bats: a natural reservoir of Hendra virus. Journal of General Virology. 2000;81(8):1927–32.

16. Freiberg A, Dolores LK, Enterlein S, Flick R. Establishment and characterization of plasmid-driven minigenome rescue systems for Nipah virus: RNA polymerase I- and T7-catalyzed generation of functional paramyxoviral RNA. Virology. 2008;370(1):33–44.

17. Halpin K, Bankamp B, Harcourt BH, Bellini WJ, Rota PA. Nipah virus conforms to the rule of six in a minigenome replication assay. Journal of General Virology. 2004;85(3):701–7.

18. Watt A, Moukambi F, Banadyga L, Groseth A, Callison J, Herwig A, et al. A Novel Life Cycle Modeling System for Ebola Virus Shows a Genome Length-Dependent Role of VP24 in Virus Infectivity. Journal of Virology. 2014-9-15;88(18).

19. Calain P, Roux L. The rule of six, a basic feature for efficient replication of Sendai virus defective interfering RNA. J Virol. 1993;67(8):4822–30.

20. Kolakofsky D, Pelet T, Garcin D, Hausmann S, Curran J, Roux L. Paramyxovirus RNA Synthesis and the Requirement for Hexamer Genome Length: the Rule of Six Revisited. Journal of Virology. 1998;72(2):891–9.

21. Ker D-S, Jenkins HT, Greive SJ, Antson AA. CryoEM structure of the Nipah virus nucleocapsid assembly. PLOS Pathogens. Jul 16, 2021;17(7).

22. Walpita P, Peters CJ, Walpita P, Peters CJ. Cis-acting elements in the antigenomic promoter of Nipah virus. Journal of General Virology. 2007/09/01;88(9).

23. Noton SL, Fearns R. INITIATION AND REGULATION OF PARAMYXOVIRUS TRANSCRIPTION AND REPLICATION. Virology. 2015/05;0.

24. Ouizougun-Oubari M, Fearns R, Ouizougun-Oubari M, Fearns R. Structures and Mechanisms of Nonsegmented, Negative-Strand RNA Virus Polymerases. Annual Review of Virology. 2023/09/29;10(Volume 10, 2023).

25. Beaty SM, Park A, Won ST, Hong P, Lyons M, Vigant F, et al. Efficient and Robust Paramyxoviridae Reverse Genetics Systems. mSphere. 2017;2(2).

26. Haas G, Lee B. Reverse Genetics Systems for the De Novo Rescue of Diverse Members of Paramyxoviridae. Methods in Molecular Biology. 2024.

27. Park A, Yun T, Hill TE, Ikegami T, Juelich TL, Smith JK, et al. Optimized P2A for reporter gene insertion into Nipah virus results in efficient ribosomal skipping and wild-type lethality. Journal of General Virology. 2016;97(4):839–43.

28. Wang YE, Park A, Lake M, Pentecost M, Torres B, Yun TE, et al. Ubiquitin-Regulated Nuclear-Cytoplasmic Trafficking of the Nipah Virus Matrix Protein Is Important for Viral Budding. PLoS Pathogens. 2010/11;6(11).

29. Pentecost M, Vashisht AA, Lester T, Voros T, Beaty SM, Park A, et al. Evidence for Ubiquitin-Regulated Nuclear and Subnuclear Trafficking among Paramyxovirinae Matrix Proteins. PLOS Pathogens. Mar 17, 2015;11(3).

30. Deffrasnes C, Marsh GA, Foo CH, Rootes CL, Gould CM, Grusovin J, et al. Genome-wide siRNA Screening at Biosafety Level 4 Reveals a Crucial Role for Fibrillarin in Henipavirus Infection. PLoS Pathogens. 2016/03;12(3).

31. Li Z, Yu M, Zhang H, Wang H-Y, Wang L-F. Improved rapid amplification of cDNA ends (RACE) for mapping both the 5′ and 3′ terminal sequences of paramyxovirus genomes. Journal of Virological Methods. 2005;130(1):154–6.

32. Sourimant J, Lieber CM, Aggarwal M, Cox RM, Wolf JD, Yoon JJ, et al. 4′-Fluorouridine is an oral antiviral that blocks respiratory syncytial virus and SARS-CoV-2 replication. Science. 2022;375(6577):161-+.

33. Cox RM, Sourimant J, Toots M, Yoon JJ, Ikegame S, Govindarajan M, et al. Orally efficacious broad-spectrum allosteric inhibitor of paramyxovirus polymerase. Nat Microbiol. 2020;5(10):1232–46.

34. Jordan PC, Liu C, Raynaud P, Lo MK, Spiropoulou CF, Symons JA, et al. Initiation, extension, and termination of RNA synthesis by a paramyxovirus polymerase. PLOS Pathogens. Feb 9, 2018;14(2).

35. Calain P, Roux L. Functional Characterisation of the Genomic and Antigenomic Promoters of Sendai Virus. Virology. 1995/09/10;212(1):163–73.

36. Pelet T, Marq J-B, Sakai Y, Wakao S, Gotoh H, Curran J, et al. Rescue of Sendai virus cDNA templates with cDNA clones expressing parainfluenza virus type 3 N, P and L proteins. Journal of General Virology. 1996/10/01;77(10).

37. Brown DD, Collins FM, Duprex WP, Baron MD, Barrett T, Rima BK, et al. ‘Rescue’ of mini-genomic constructs and viruses by combinations of morbillivirus N, P and L proteins. Journal of General Virology. 2005/04/01;86(4).

38. Magoffin DE, Mackenzie JS, Wang LF. Genetic analysis of J-virus and Beilong virus using minireplicons. Virology. 2007;364(1):103–11.

39. Yun T, Park A, Hill TE, Pernet O, Beaty SM, Juelich TL, et al. Efficient reverse genetics reveals genetic determinants of budding and fusogenic differences between Nipah and Hendra viruses and enables real-time monitoring of viral spread in small animal models of henipavirus infection. J Virol. 2015;89(2):1242–53.

40. Garner OB, Yun T, Pernet O, Aguilar HC, Park A, Bowden TA, et al. Timing of Galectin-1 Exposure Differentially Modulates Nipah Virus Entry and Syncytium Formation in Endothelial Cells. Journal of Virology. 2015/03/03;89(5).

